# A novel Golgi-associated Vangl2 translational variant required for PCP regulation in vertebrates

**DOI:** 10.1101/2020.10.20.347385

**Authors:** Alexandra Walton, Diego Revinski, Arnauld Sergé, Stéphane Audebert, Luc Camoin, Tania M Puvirajesinghe, Daniel Isnardon, Sylvie Marchetto, Laurent Kodjabachian, Eric Bailly, Jean-Paul Borg

## Abstract

First described in *Drosophila melanogaster*, planar cell polarity (PCP) is a developmental process essential for embryogenesis and development of polarized structures in Metazoans. This signaling pathway involves a set of evolutionarily conserved genes encoding transmembrane (Vangl, Frizzled, Celsr) and cytoplasmic (Prickle, Dishevelled) molecules. Vangl2 is of major importance in embryonic development as illustrated by its pivotal role during neural tube closure in human, mouse, *Xenopus* and zebrafish embryos. The regulated and poorly understood traffic of Vangl2 to the plasma membrane is a key event for its function in development. Here we report on the molecular and functional characterization of a novel 569-amino acid N-terminally extended Vangl2 isoform, Vangl2-Long, that arises from an alternative non-AUG translation initiation site, lying 144 base pair upstream of the conventional start codon. While missing in Vangl1 paralogs and in all invertebrates, including *Drosophila melanogaster*, this N-terminal extension is conserved in all vertebrate Vangl2 sequences and confers a subcellular localization in the Golgi apparatus, probably as a result of an extended retention time in this organelle. Vangl2-Long belongs to a multimeric complex with Vangl1 and Vangl2 and we show that its down-regulation leads to severe PCP-related phenotypes in *Xenopus* embryos, including shorter body axis and neural tube closure defects. Altogether, our study unveils a novel level of complexity in Vangl2 expression, trafficking and function.

## Introduction

Planar cell polarity (PCP) refers to the cell polarization within the plane of the epithelial sheet and is crucial for the regulation of embryonic development and tissue formation (1). Mammalian Vangl2, known as Van Gogh (Vang)/Strabismus (Stbm) in the fruit fly *Drosophila melanogaster*, is a member of the core PCP proteins, which also comprise Celsr, Frizzled, Dishevelled and Prickle. First discovered in the fly as a regulator of eye, wing and bristle development (2,3), Vang/Stbm is highly conserved during evolution. Like Vang/Stbm, the vertebrate orthologs Vangl1 and Vangl2 (4,5) contain four transmembrane domains with cytoplasmic amino- and carboxy-termini and two short loops facing the extracellular space (6–8). In addition, their carboxy-terminal region ends with a conserved PDZ binding motif (PBM) (3,9). Previous studies have established that the two paralogs interact both genetically (10) and physically (11), with Vangl2 being able to homodimerize and to form heterodimers with Vangl1.

Loss of function mutations in human *Vangl1* and *Vangl2* genes are associated with neural tube defects (NTDs) (12,13), highlighting the importance of these genes in developmental processes. Similarly, mouse mutants with a characteristic *Loop-tail (Lp)* phenotype are deficient for Vangl2 functions and *Lp/Lp* homozygous mice display the most severe failure in neural tube closure (craniorachischisis) (14). NTDs result from defective convergent-extension (CE) movements, a collective and polarized migration of mesenchymal cells during gastrulation and neurulation stage. In *Xenopus* and zebrafish, disrupted CE movements during gastrulation and neurulation due to the depletion of Vangl2 are respectively associated with a severe reduction in the body length (15,16) and incomplete neural tube closure (17–19).

Like Vang/Stbm, Vangl2 localizes at the plasma membrane (PM) in planar polarized epithelial tissues (20–22). Recent findings have highlighted that the traffic of Vangl2 to the PM is essential for proper development, as Vangl2 function mostly relies on its cellular localization. The molecular components involved in Vangl2 trafficking to the surface are progressively emerging (23). Sec24b, a component of the coat protein complex II (COPII), has been shown to play a key role in the sorting of Vangl2 from the endoplasmic reticulum (ER). Similar to *Vangl2 Lp* mutant, loss-of-function mutation in *Sec24b* leads to mislocalization of Vangl2 in the ER and consequently to neural tube defects (24,25) (25,26). The transport of Vangl2 from the trans-Golgi network (TGN) to the cell surface has also been investigated. Guo et al. provided evidence that the small GTPase Arfrp1, together with the clathrin adaptor protein 1 (AP-1), regulates the export of Vangl2 from the TGN (27,28). They identified an YYXXF TGN sorting signal in the intracellular carboxy-terminal region of Vangl2. Once localized at the PM Vangl2 is probably endocytosed, yet the mechanisms controlling this step are poorly described. Recent studies have proposed that Vangl2 endocytosis is dependent on dynamin and Rab5 (29) and that the trafficking regulatory factor GIPC1, which contains a PDZ domain that binds to the PDZ-binding motif (PBM) of Vangl2, increases the localization of Vangl2 in endocytic vesicles (30). Recycling of endocytosed Vangl2 from endosomes to the PM or to the TGN could involve the early endosome-associated retromer component SNX27 (31), which also contains a PDZ domain that binds to the PBM of Vangl2 (32).

While most eukaryotic translation initiation events are known to take place at canonical AUG start codons present near the 5’ cap of messenger RNAs, increasing numbers of translation initiation events have been reported to take place at near-cognate start codons (33–36). Because these alternative translation initiation sites (ATIS) are often found in the 5’-UTR cap region of mRNAs and in-frame with the downstream canonical AUG start codon, their usage frequently yields longer isoforms owing to the presence of additional N-terminal sequences that may provide them with distinct biological functions relative to AUG-initiated isoforms (37–47). Alternative translation initiation is therefore recognized as an additional source of isoform diversity.

Here we identify a novel N-terminally extended isoform of Vangl2, termed Vangl2-Long, that arises from the use of a non-AUG start codon upstream of the coding region of canonical Vangl2. Vangl2-Long contains an evolutionarily conserved N-terminal 48 amino acid sequence bearing a signal for subcellular localization in the Golgi apparatus. Moreover, we provide data showing that Vangl2-Long is important for convergent extension and neural tube closure in *Xenopus.* These data describe a further level of complexity in Vangl2 expression, trafficking and function.

## Results

### Identification of a novel N-terminally extended human and murine Vangl2 isoform

Biochemical characterization of a monoclonal antibody (mAb 36E3) raised against the N-terminal region of human Vangl2 (11) revealed that, in addition to the main 62 kDa band corresponding to the 521aa long Vangl2 polypeptide, this antibody also detected an antigen with an apparent molecular weight of ≈70kDa, in both human and murine cells (Figure 1A). The additional 70kDa band was systematically present in all Vangl2 positive cell lines and of lower intensity compared to the major 62kDa Vangl2 signal (Figure 1A, 1G). Another previously characterized Vangl2 antibody, mAb 2G4, similarly detected two bands by Western blot in mouse tissues (11). *Vangl2* knockout with a CRISPR-Cas9 approach abrogated the expression of both the 62 and 70 kDa antigens (Figure S1A). Taken together these observations made the hypothesis of a spurious cross-reactivity between the two bands highly unlikely and led us to explore the possibility of a new Vangl2 isoform. While protein isoforms can arise from many different molecular events including post-translational modifications, alternative splicing or alternative translation initiation, we focused our attention on the latter scenario for two reasons. First, previous mass spectrometry analyses of human Vangl2 (11) identified a peptide harboring an extra three amino acid (SDA) sequence upstream of the initiating methionine (Figure 1C), evoking the possibility of an N-terminally extended isoform. Second and also fully consistent with this scenario, an *in-silico* study suggested the presence of a near-cognate translation initiation AUA codon lying 144 nucleotides upstream of the canonical AUG initiation codon of Vangl2 (Figure 1B) (33). Because this alternative AUA initiation site is in-frame with the canonical AUG start codon and carried by the same exon, it was expected to add a 48aa N-terminal extension to the conventional 521aa Vangl2 protein (Figure 1B and 1C), potentially yielding a 569aa Vangl2 isoform (hereafter referred to as Vangl2-Long) with a calculated MW of 65 kDa, close to the size of the observed 70 kDa band.

**Figure 1.**
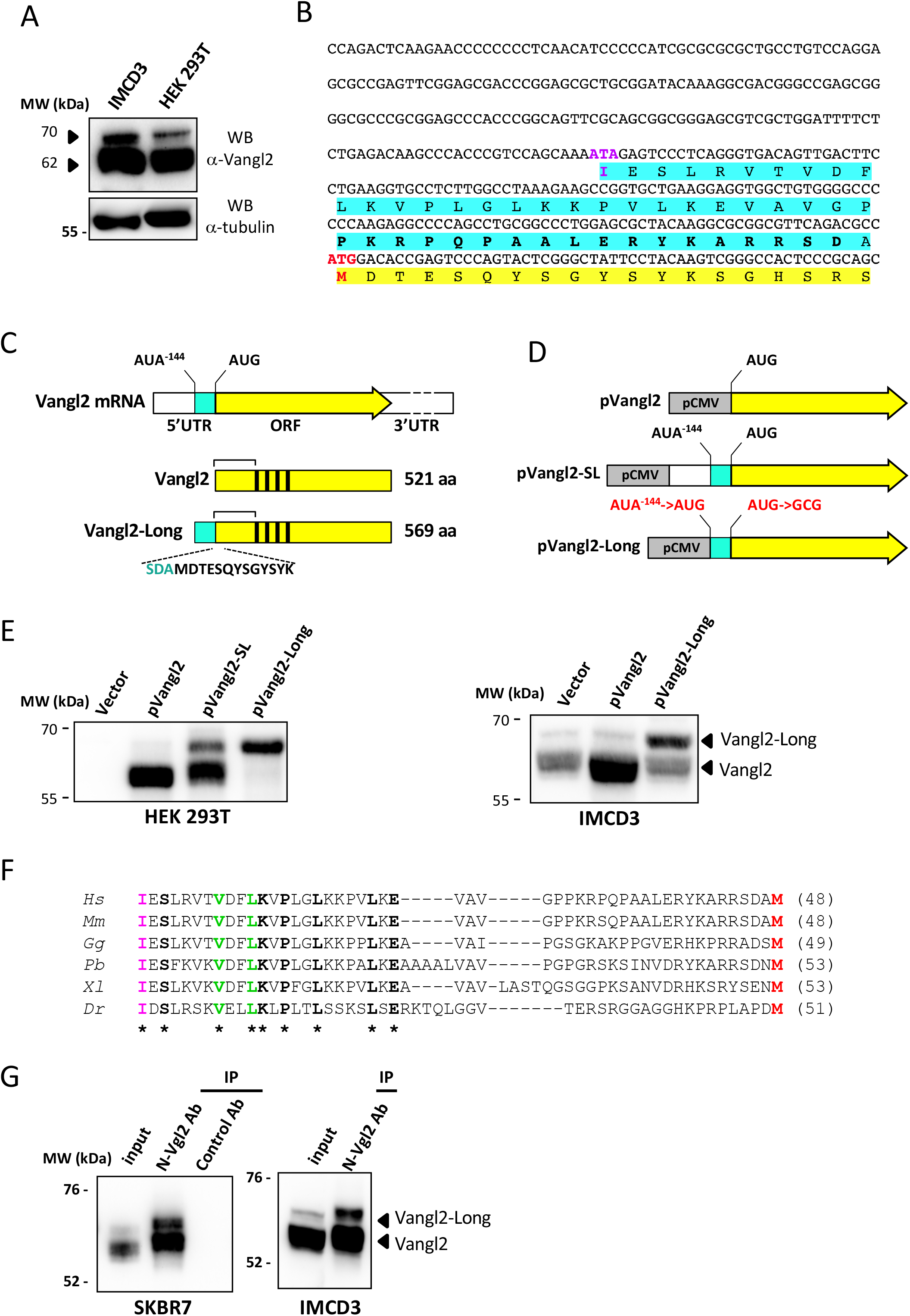
Identification of Vangl2-Long, a novel highly-conserved N-terminally extended Vangl2 isoform, in human and mouse cells. **(A)** Western blot analysis of murine epithelial (IMCD3) and human embryonic kidney (HEK 293T) cell extracts using Vangl2-specific monoclonal antibody mAb 36E3 (top panel). Arrows point to the major 62 KDa Vangl2 protein and to a less intense and slower migrating band of 70 KDa both recognized by mAb 36E3. Immunoblot with an anti-alpha tubulin (botton panel) was used as a loading control **(B)** DNA sequence of a 5’ region of human Vangl2 cDNA encompassing 342bp of the 5’UTR region immediately upstream to the conventional ATG start site (highlighted in red) and the first 57bp of ATG-initiated Vangl2 ORF. A potential ATA alternative initiation site in position - 144 (ATA^-144^) relative to the canonical Vangl2 ATG^1^ start site is highlighted in magenta. Amino acid sequences of Vangl2 and the N-terminal extension of Vangl2-Long are highlighted in yellow and cyan, respectively. Sequence of the peptide used to generate the N-Vgl2 pAb rabbit antibodies is indicated in bold letters **(C)** Schematic representations of the human Vangl2 mRNA (top) with its canonical (AUG) and near-cognate (ATA^-144^) initiation sites and the two Vangl2 (middle) and Vangl2-Long (bottom) encoded isoforms. Met-initiated Vangl2 and the ATA-initiated N-terminal extension are drawn in yellow and cyan, respectively. Black rectangles represent the four transmembrane domains present in both isoforms and numbers on the right give their respective length. Sequence of a peptide identified by mass spectrometry and encompassing the N-terminus of Vangl2 (black) preceded by 3 residues of the 5’UTR region (cyan) is shown below Vangl2-Long. Brackets above the two schematized isoforms indicate a common region recognized by mAb 36E3 **(D)** Schemes of pVangl2 (top), pVangl2-SL (middle) and pVangl2-Long (bottom) vectors. All Vangl2 constructs are under the control of CMV promoter (grey box) and Vangl2 isoform representation uses the same color code as in (C). Note that, as indicated in red in pVangl2-Long, AUA ^-144^ and the canonical AUG codon have been mutated to, respectively, AUG and GCG in order to enhance Vangl2-Long biosynthesis while abrogating Vangl2 expression by this construct. **(E)** HEK 293T (left panel) and IMCD3 (right panel) cells were probed by immunoblotting with anti-Vangl2 mAb 36E3 upon transfection with the indicated plasmids. Note that the absence of endogenous Vangl2 signals in HEK 293T cells transfected with an empty plasmid (vector) is due to the short exposure time used to detect the overexpressed Vangl2 isoforms. Arrows point to Vangl2 and Vangl2-Long isoforms that are overexpressed by the relevant pVangl2 and pVangl2-Long constructs. **(F)** Amino acid sequence alignment of predicted N-terminal extensions from six vertebrate species, including human *(Hs)*, mouse (Ms), chicken *(Gg)*, python *(Pb)*, African clawed frog *(Xl)* and zebrafish *(Dr).* Residues conserved in the extension of all six species are indicated in bold letters and by asterisks. AUA-encoded isoleucine residues that initiate the translation of all the extensions shown here are highlighted in magenta whereas the AUG-encoded methionine residue of Vangl2 is in red. Two highly conserved residues (V8 and L11) whose mutagenesis is described in a later section of this study are highlighted in green. **(G)** Proteins extracted from SKBR7 (left panel) and IMCD3 (right panel) cells were immunoprecipitated with the Vangl2-Long isoform-specific N-Vgl2 pAb antibodies and immunopurified proteins were immunoblotted with mAb 36E3. Control pAb is an isotypic control rabbit antibody.

To address whether the 5’UTR region of *Vangl2* mRNA can direct the translation of a second and longer isoform, as predicted above, we designed three expression plasmids (Figure 1D). In the first construct (pVangl2), Vangl2 ORF initiated by the canonical AUG start codon was cloned downstream of the strong CMV promoter. A second plasmid (pVangl2-SL) was a 5’ extended version of pVangl2 in which the 342pb region immediately upstream of the Vangl2 ORF (Figure 1D) was inserted in frame with the AUG start codon of Vangl2. A third vector (pVangl2-Long) was specifically generated for the expression of a 569aa Vangl2-Long polypeptide, using an appropriately designed cDNA as depicted in Figure 1D. Upon transfection in HEK 293T and IMCD3 cells, pVangl2 and pVangl2-Long plasmids each drove the synthesis of a single polypeptide of, respectively, 62 and 70 kDa, that perfectly comigrated with endogenous Vangl2 antigens (Figure 1E), therefore indicating that the 70 kDa band likely corresponds to Vangl2-Long. Remarkably, in the same transfection assay, pVangl2-SL simultaneously gave rise to two distinct 62 and 70 kDa Vangl2 products with a Vangl2:Vangl2-Long ratio that mirrored the ratio observed in mammalian cell extracts (compare left panel of Figure 1E to Figure 1A). This result fully established the ability of the 5’UTR mRNA to initiate the translation of a longer and less abundant Vangl2 isoform, in addition to the major canonical Vangl2 protein. Consistent with a critical role of the upstream AUA codon, an ATA-to-GCA mutation at position 144 in pVangl2-SL abrogated the expression of Vangl2-LONG without altering the levels of Vangl2 (Figure S2, ATA^-144^->GCA lane). Taken together, these data provide strong experimental evidence for the requirement of AUA^144^ in the alternative translation initiation of a novel Vangl2 isoform, that differs from the canonical isoform by the presence of a 48 aa N-terminal extension.

The ability of the *Vangl2* locus to encode a longer isoform from a non-canonical AUA start codon is not limited to human but found in all vertebrates for which a Vangl2 sequence is currently available. In contrast, this N-terminal extension is found neither in the Vangl1 mRNA nor in the orthologous protein of more distantly related species such as *Drosophila melanogaster* or *Caenorhabditis elegans.* Protein sequence comparison between the N-terminal extensions of mammals, birds, reptiles, amphibians and fishes confirmed the high degree of phylogenetic conservation of this motif (Figure 1F). This is best illustrated by the presence of several invariant residues along the extension of all these vertebrate members. The protein sequence of the 48aa Vangl2-Long extension has however no homology with any other sequences available in protein databases.

We next sought to validate the expression and identity of Vangl2-Long in mammalian cell lines, using rabbit polyclonal antibodies (hereafter referred to as N-VGL2 pAb) raised against a peptide present within the N-terminal extension of Vangl2-Long (Figure 1B). An immunoprecipitation (IP) assay with IMCD3 cells engineered to express either of the two Vangl2 isoforms confirmed the ability of N-VGL2 pAb to recognize Vangl2-Long but not Vangl2 (Figure S1C). Further IP experiments with this tool allowed us to demonstrate the occurrence of endogenous Vangl2-Long in human (SKBR7) and murine (IMCD3) cells (Figure 1G) as evidenced by the strong enrichment of a 70kDa 36E3-reactive signal in the N-VGL2 pAb IP. Surprisingly, the 62 kDa Vangl2 protein was also massively detected in the material immmunoprecipitated by N-VGL2 pAb and not with a control antibody (Figure 1G). Since N-VGL2 pAb does not recognize Vangl2 alone (Figure S1C), the latter observation led us to conclude to the existence of a physical interaction between Vangl2 and Vangl2-Long. Taken together the biochemical data described above clearly identify the 70 kDa protein as a naturally occurring Vangl2-Long isoform in human and murine cells and establish its ability to form an endogenous complex with Vangl2.

### Vangl2 isoforms associate to Vangl1 in a multimeric complex

The two paralogs Vangl1 and Vangl2 are known to interact both genetically (10) as well as physically (11). To assess whether Vangl2-Long can similarly hetero-oligomerize with Vangl1, lysates of HEK 293T cells expressing GFP-Vangl1 alone or in combination with Vangl2 or Vangl2-Long were immunoprecipitated with anti-Vangl2 mAb 36E3 or the Vangl2-Long specific antibody N-VGL2 pAb (Figure 2A) and monitored by WB for Vangl2 and GFP-Vangl1. This experiment confirmed the previous observation that Vangl1 and Vangl2 form a complex. It also showed that GFP-Vangl1 co-purifies with Vangl2-Long regardless of the antibody used for the IP, demonstrating that that Vangl2-Long, like Vangl2, is able to interact with Vangl1.

**Figure 2.**
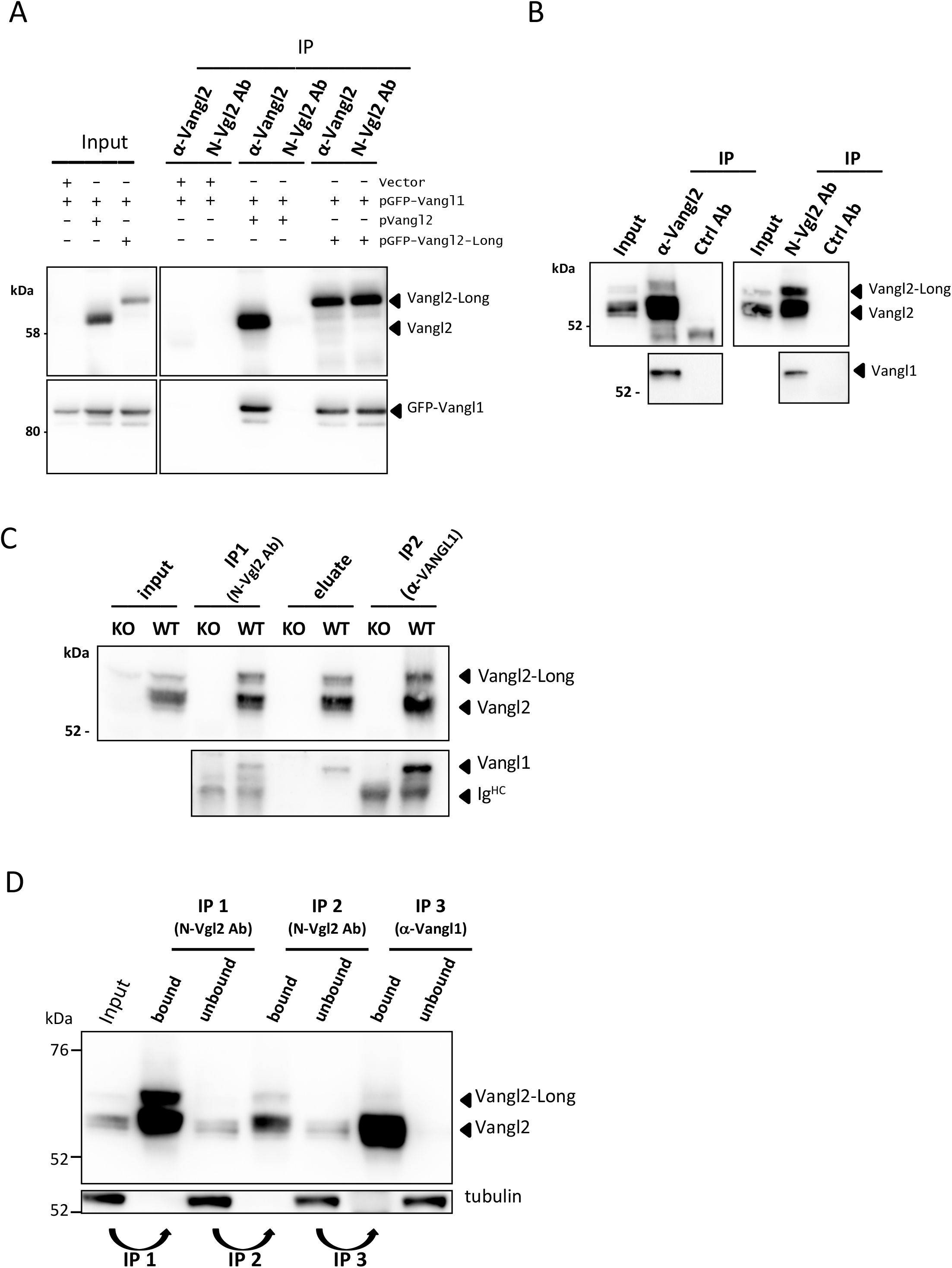
Vangl2-Long forms a tri-partite complex with Vangl1 and Vangl2 in vivo. **(A)** Immunoprecipitation of lysates from HEK 293T cells transfected with GFP-Vangl1 and a control plasmid or GFP-Vangl1 in combination with pVangl2 or pVangl2Long, using mAb 36E3 or N-Vgl2 pAb antibodies, as indicated. The presence of GFP-Vangl1 and Vangl2 in the immunoprecipitate samples was determined by Western blotting using, respectively, a GFP-specific antibody (bottom panels) and mAb 36E3 (top panels). Arrows point to the Vangl2 and Vangl2-Long signals detected by mAb 36E3 **(B)** SKBR7 cell extracts were immunoprecipitated with either mAb 36E3 (left panels) or N-VGL2 pAb (right panels) followed by a Western blot analysis of Vangl1 (bottom panels) and Vangl2 isoforms (top panels) using, respectively mAb 19D5 and mAb 36E3. Isotopic rat and rabbit antibodies were used as negative controls for each IP experiments (Ctrl Ab). **(C)** Lysates from wild type (WT) and Crispr-Cas9 KO-VANGL2 (KO) HEK 293T cells were first immunoprecipitated with N-VGL2 pAb (IP1, N-Vgl2 Ab). Bound proteins were subsequently eluted with the relevant immunogenic peptide (eluate) before being subjected to a second round of IP with Vangl1-specific mAb 19D5 (IP2, α-Vangl1). The different fractions were examined by immunoblotting for their Vangl1 (lower panel) and Vangl2 (upper panel) contents using, respectively, mAb 19D5 and mAb 36E3. Arrows in the upper panel indicate the two endogenous Vangl2 and Vangl2-Long isoforms while those in the lower panel point to Vangl1 and immunoglobulin heavy chains (lg^HC^). **(D)** Biochemical evidence for the existence of binary and ternary complexes between Vangl1, Vangl2 and Vangl2-Long in IMCD3 cells. An IMCD3 cell extract was immunoprecipitated twice with N-Vgl2 pAb (IP1, and IP2, N-Vgl2-Ab) in order to deplete Vangl2-long before subjecting the unbound proteins to a third immunoprecipitation step using a Vangl1 specific antibody, mAb 3D1 (IP3, α-Vangl1). Immunoprecipitates (bound) and IP supernatants (unbound) were probed for the presence of Vangl2 and Vangl2-Long by Western blotting with mAb 36E3. An α-tubulin immunoblot was performed in parallel to serve as a protein loading control. Note the strong enrichment of Vangl2 and Vangl2-Long in the IP1 sample and the extensive depletion of Vangl2-Long after the second round of immunoprecipitation with N-Vgl2 pAb (unbound, IP2). Also note that a pool of Vangl2 not immunoprecipitated by N-Vgl2 pAb efficiently co-purifies with Vangl1 in the α–Vangl1 IP3 fraction (bound, IP3)

We next asked if complexes made of Vangl1 and either of the two Vangl2 isoforms could be detected *in vivo* in SKBR7 cells as this cell line is known to express all of these VANGL proteins. To address this issue, we took advantage of recently developed Vangl1-specific mAbs (Figure S3) to monitor the presence of Vangl1 among the proteins immunoprecipitated from SKBR7 cell lysates with either mAb 36E3 or N-VGL2 pAb antibodies. In both cases Vangl1 was readily co-immunoprecipitated (Figure 2B) highlighting the capacity of Vangl1 to form a protein complex with the Vangl2 and Vangl2-Long isoforms in SKKBR7 cells.

The biochemical data presented above point to a unique property of the three Vangl1/2/2-Long proteins to physically interact with one another *in vivo.* Yet, the question remained as to whether these proteins can assemble into a single macromolecular complex. We addressed this question by performing a sequential immunoprecipitation assay in which the Vangl2-Long isoform and its associated proteins were first immunopurified from HEK 293T cells with N-VGL2 pAb (Figure 2C, IP1 N-Vgl2 Ab). A Vangl2-KO HEK 293T cell line was used as a negative control for this experiment. Proteins retained by the N-Vgl2 pAb antibody were subsequently eluted under native conditions and subjected to a second round of immunoprecipitation with a Vangl1 antibody (Figure 2C, IP2 α-Vangl1). Western blot analysis was then conducted to monitor Vangl1 and Vangl2 contents of the different fractions. As expected from data illustrated in Figures 1G and 2B, immunoprecipitation with N-VGL2 pAb resulted in a substantial enrichment of VANGL2-Long but also Vangl2 and Vangl1 (Figure 2C, IP1). Importantly, all these proteins were successfully eluted from the beads as judged by the identical Vangl1 and Vangl2/2-Long profiles observed in the N-VGL2 pAb IP and eluate fractions (compare fourth and sixth lanes of Figure 2C). The presence of both Vangl2 and Vangl2-Long isoforms in the final Vangl1 IP unambiguously established the existence of macromolecular complexes containing the three Vangl proteins (Figure 2C, IP2). These biochemical experiments also revealed that Vangl2-Long could be efficiently depleted from IMCD3 cell lysates after two successive rounds of IPs with the same N-Vgl2 pAb antibody. In contrast a significant fraction of Vangl2 was still present in the unbound materials (Figure 2D, third and fourth lanes, IP1-N-Vgl2 pAb-unbound and IP2-N-Vgl2 pAb-unbound). To determine if Vangl2 present in the unbound material was associated with Vangl1, we subjected this fraction to a third IP with a Vangl1 specific antibody. Vangl2 was not only recovered in the Vangl1 IP but also fully depleted in the material not retained by the Vangl1 antibody (Figure 2D, two right lanes, IP3-αVangl1-bound and unbound fractions) providing evidence for a second pool of Vangl2 associated with Vangl1. Altogether these data suggest the existence of at least two types of Vangl oligomers in human and mouse cells: a tripartite complex that include Vangl1, Vangl2 and Vangl2-Long and a simpler complex made of Vangl1 and Vangl2 only.

### isoform-specific localization of VangL2 in IMCD3 cells

We next sought to compare the cellular distribution of the two Vangl2 isoforms in IMCD3 cells. We chose this cell line as it constitutes a convenient epithelial cell polarity model forming extensive intercellular junctions when cells are grown to high confluency. Because N-VGL2 pAb is not suitable for immunofluorescence experiments, we analyzed by immunofluorescence (IF) microscopy IMCD3 cells stably transfected with GFP-Vangl2 or GFP-Vangl2-Long constructs. Examination of GFP-Vangl2 distribution in confluent IMCD3 monolayers confirmed previous results (19) as most of the protein was observed at E-cadherin-decorated cell–cell contacts (Figure 3A, upper panels and Figure 3B). This was in marked contrast with GFP-Vangl2-Long whose plasma membrane recruitment was significantly reduced and which was instead found associated with internal membrane compartments including the *cis*-Golgi compartment (Figure 3A, lower panels and Figure 3B) and the *trans*-Golgi network (TGN) (Figure S4).

**Figure 3.**
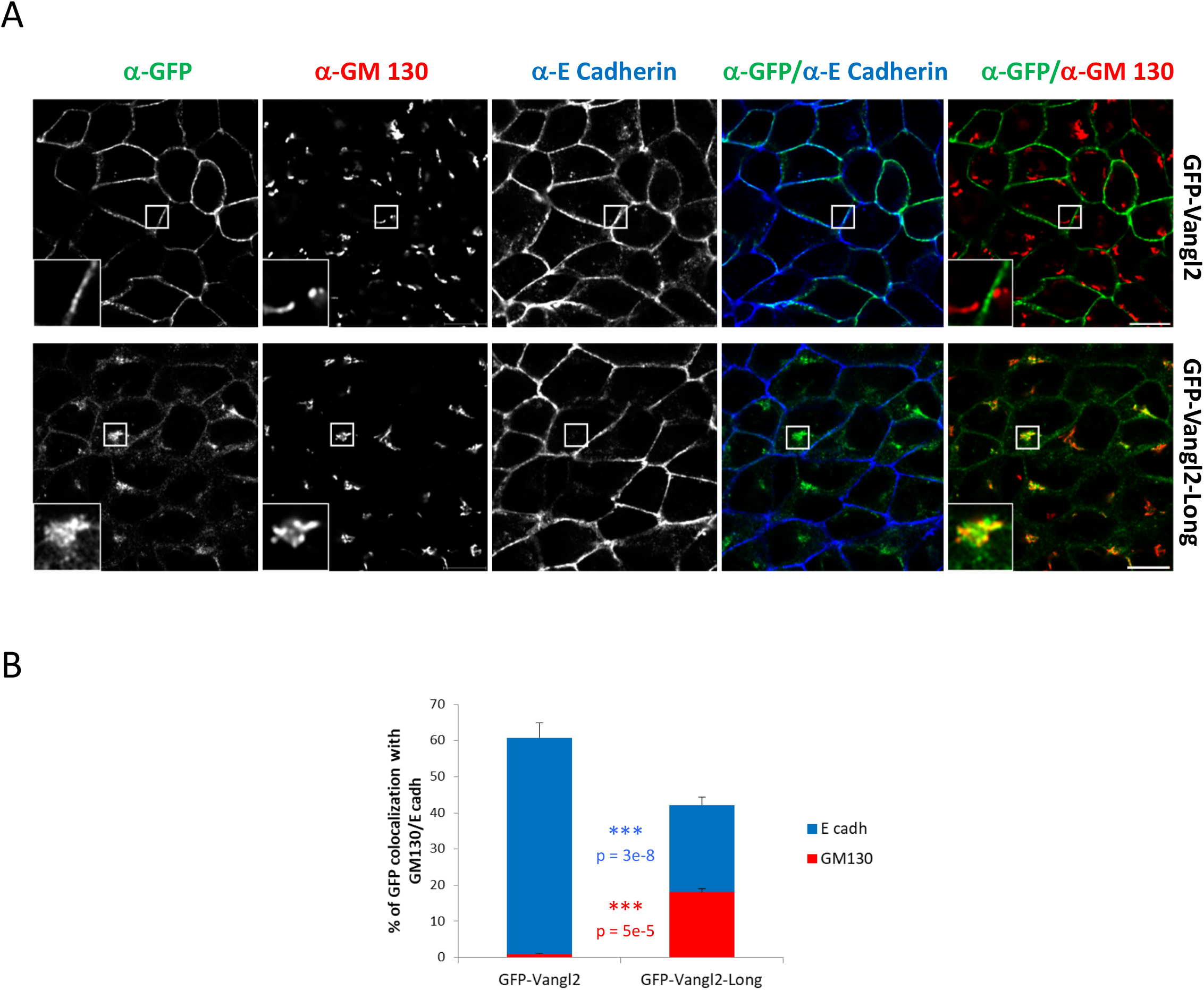
Cellular localization of GFP-tagged Vangl2 and Vangl2-Long isoforms in confluent IMCD3 cells. **(A)** IMCD3 cells stably transfected with pGFP-Vangl2 or pGFP-Vangl2-Long constructs were grown to high confluency before being processed for immunofluorescence microscopy with the indicated antibodies. Scale bar represents 10 μm. **(B)** Quantification of the percentage of GFP-Vangl2 and GFP-Vangl2-Long co-localizing with E-Cadherin (blue) and GM130 (red) fluorescence signals using MATLAB code. Histograms represent the mean ± s.e.m calculated for three independent experiments (>30 cells each). P-values evaluated by Student’s t-test. *** p<1e-4.

A very similar picture was obtained when assessing the localization of Vangl2 and Vangl2-long proteins in IMCD3 Vangl2 KO cells, whether these proteins were GFP-tagged or not (Figures S5 and S6). These observations indicate that the GFP tag has no impact on the cellular localization of the Vangl2 isoforms. The finding that the localization pattern was unchanged in KO Vangl2 cells also suggests that the localization of these isoforms is largely independent of each other. We also observed that under low confluency conditions, GFP-Vangl2 exhibited an increased intracellular distribution (Figure S7B upper panels). In agreement with the above results, GFP-Vangl2-Long was also frequently associated with the Golgi apparatus (Figure S7B, lower panels). Taken together, these results unveil the enhanced propensity of the newly identified Vangl2-Long isoform to associate with the Golgi compartment.

### Vangl2-Long has a longer retention time in the Golgi apparatus

Given the enhanced propensity of Vangl2-Long to locate at the Golgi apparatus when compared to Vangl2, we wondered if these two isoforms transit similarly through this organelle via the canonical endoplasmic reticulum (ER)-Golgi secretory pathway. Recent findings have highlighted the importance of proper sorting of Vangl2 from the ER and the TGN compartments (25,27). Yet, vesicular trafficking of Vangl2 from the ER to the cis-Golgi compartment has never been investigated. To explore the dynamics of Vangl2 isoform trafficking, we took advantage of the Retention Using Selective Hooks (RUSH) assay which enables to study and quantify the synchronous transport of cargo molecules in living cells (53). Accordingly, Vangl2 and Vangl2-Long were cloned downstream of a Streptavidin Binding Peptide (SBP)-GFP cassette in a RUSH plasmid that also expressed an ER-targeted hook constituted by the streptavidin-Ii fusion protein (53). When transfected in IMCD3 cells, the resulting RUSH-Vangl2 and RUSH-Vangl2-Long plasmids directed the expression of similar protein levels (Figure S8A). In this experimental setting, the Strep-Ii fusion hook protein efficiently sequestered both Vangl2 isoforms in the ER, in the absence of exogenous biotin (Figure S8B) while biotin addition led, as expected, to their rapid exit from this compartment (data not shown, see below). To help visualize the Golgi apparatus during biotin-induced trafficking of Vangl2 isoforms, cells were co-transfected using a vector encoding N-acetylgalactosaminyltransferase (GalNac-T), a human Golgi resident enzyme fused to RFP. Real-time imaging performed just before biotin treatment revealed a punctate localization pattern of both Vangl2 isoforms (Figure 4A, 4B t=0 min) that was fully consistent with the ER retention documented by our IF data (Figure S8B). By 15 min after biotin addition, the Vangl2 cargo appeared to have entirely relocated from the ER to the Golgi compartment where it perfectly colocalized with RFP-GalNac-T (Figure 4A, t=15 min). Likewise, Vangl2-Long rapidly left the ER after biotin addition and reached the Golgi compartment with a very similar kinetics (compare t=15 min panels in Figures 4A and 4B). These results therefore unambiguously demonstrate the capacity of both Vangl2 isoforms to transit through the Golgi apparatus following their release from the ER compartment.

**Figure 4.**
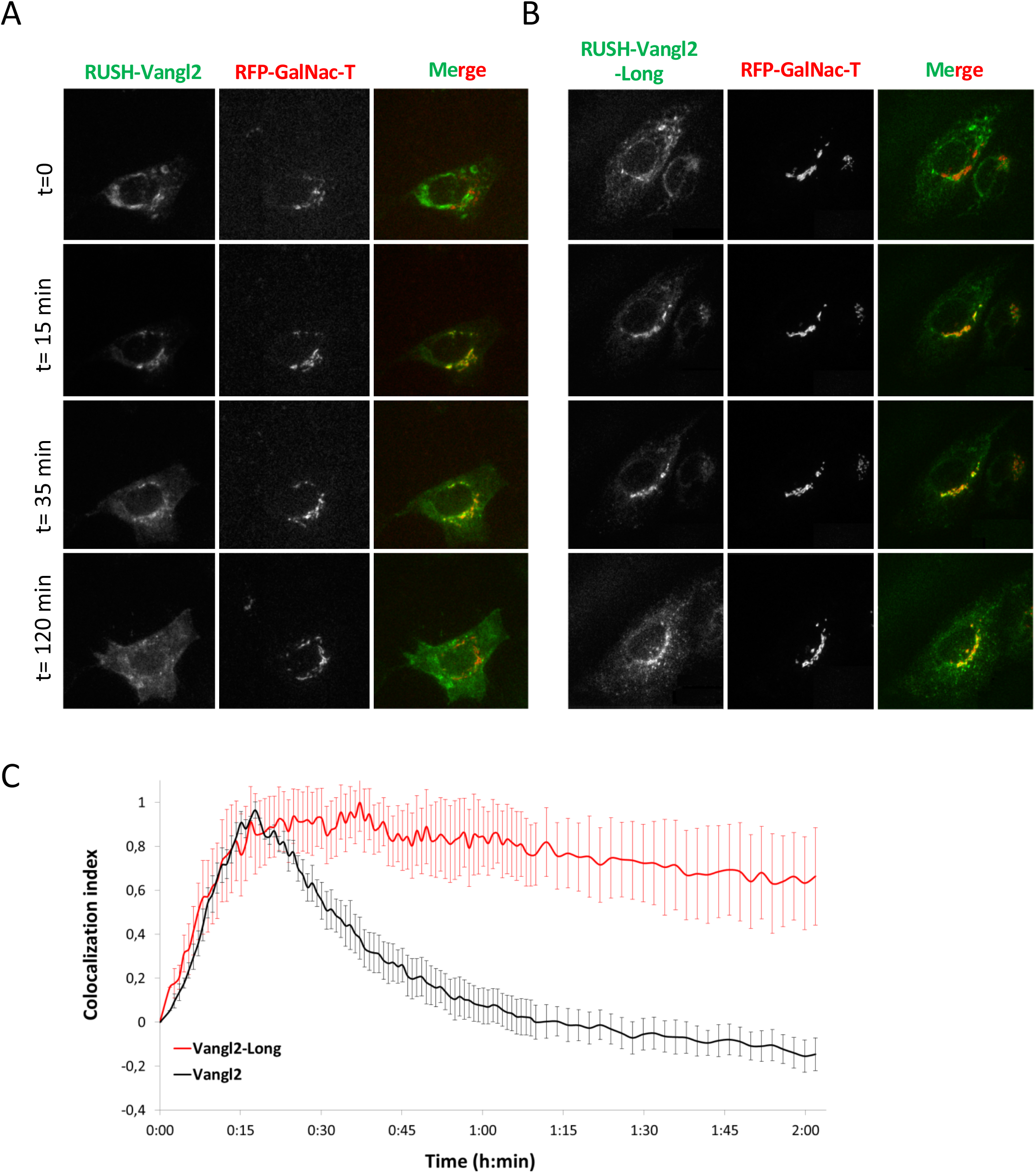
Vangl2-Long isoform shows a delayed Golgi trafficking. **(A).** Fluorescence photomicrographs of IMCD3 cells co-expressing a GFP-VANGL2 RUSH reporter and RFP-GalNac-T taken at the indicated time points before and after biotin addition. **(B)** Fluorescence photomicrographs of IMCD3 cells co-expressing a GFP-VANGL2-Long RUSH reporter and RFP-GalNac-T taken at the time points as shown in (A). **(C)** Quantification of colocalization index (mean +/− s.e.m. normalized to 0 at time 0) of GFP-VANGL2 (black) and GFP-Vangl2-Long (red) RUSH reporter signals with RFP-GalNac-T over time. n=5 videos for each condition.

Examination at later time points, however, revealed a major difference in the vesicular trafficking of the two isoforms as Vangl2 accumulated only transiently in RFP-GalNac-T-decorated Golgi stacks while the Golgi localization of Vangl2-Long persisted for a much longer period of time (compare t=35min and t=120min panels between Figure 4A and 4B). This difference was better documented by recording the co-localization index of each isoform with RFP-GalNac-T over time. As illustrated in Figure 4C, very low amounts of Vangl2 signals were present in RFP-GalNac-T-associated structures by 75 min after biotin addition while >75% of the Vangl2-Long signal was still detectable in Golgi structures by this time. Taken together, these results not only identify the Golgi apparatus as an intermediate step in the vesicular trafficking of both Vangl2 isoforms but also suggest that the Vangl2-Long isoform transits through this organelle significantly more slowly than Vangl2 and/or is actively retained in this compartment during its anterograde transport.

### The N-terminal extension of Vangl2-Long contains a Golgi-retention signal

The data presented above establish that a larger fraction of the long Vangl2 isoform resides in the Golgi compartment relative to Vangl2 (Figure 3). To determine whether this feature is specified only by the N-terminal extension, we investigated the effects of fusing the 48aa extension with the N-terminus of its closest relative, Vangl1 (Figure 5A). IMCD3 cells stably expressing GFP-Vangl1 or the resulting GFP-tagged Vangl1-long chimeric protein were monitored by Western blot (Figure 5B) and IF microscopy (Figure 5C) for the localization of the respective GFP-tagged proteins. Consistent with previous studies (8,11), GFP-Vangl1 was predominantly found at the plasma membrane of highly confluent IMCD3 cells where, as observed for GFP-Vangl2, it also extensively co-localized with E-cadherin (Figure 5C, GFP-Vangl1 panels). In contrast, the chimeric protein GFP-Vangl1-Long was less tightly recruited to the plasma membrane and instead exhibited an intracellular staining that partially overlapped with the GM130 Golgi staining (Figure 5C, GFP-Vangl1-Long panels and Figure 5D) in a manner that was highly reminiscent of the cellular distribution of GFP-Vangl2-Long (Figure 3A). These data therefore unambiguously demonstrate the ability of the 48 amino acid extension present at the N-terminus of Vangl2-Long to act as a transferable signal that promotes the accumulation of this isoform in the Golgi compartment.

**Figure 5.**
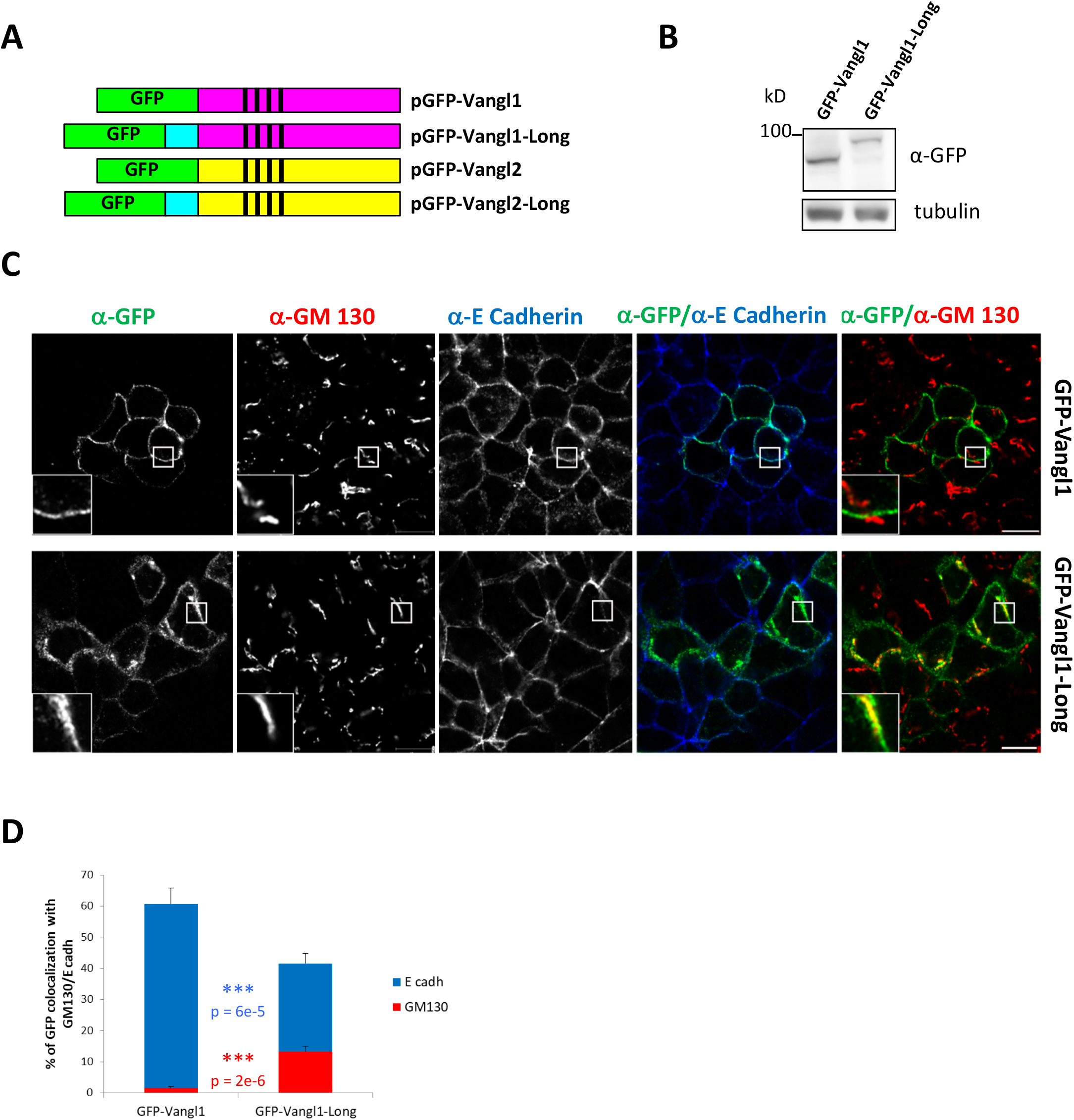
Vangl2-Long N-terminal extension acts as an autonomous Golgi-localisation signal when fused to Vangl1. **(A)** Schematic representation of Vangl1 (magenta) and Vangl2 (yellow) proteins together with their chimeric (Vangl1-Long) or naturally occurring (Vangl2-Long) N-terminally extended isoforms. The N-terminal extension region found in Vangl2-Long is indicated in cyan while green rectangles represent the GFP tags added at the aminoterminus of all Vangl1 and Vangl2 isoforms **(B)** Western blot analysis of IMCD3 cells stably expressing GFP-Vangl1 or GFP-Vangl1-Long and probed with anti-GFP (upper panel) and antitubulin (lower panel) antibodies. **(C)** Comparative immunofluorescence microscopy analysis of IMCD3 cells stably transfected with GFP-Vangl1 and GFP-Vangl1-Long. Scale bars represent 10 μm. **(D)** Quantification of the % of GFP-Vangl1 and GFP-Vangl1-Long colocalizing with E-Cadherin (blue) and GM130 (red) fluorescence signals using MATLAB code. Histograms represent the mean ± s.e.m calculated for three independent experiments (>30 cells each). P-values evaluated by Student’s t-test. *** p<1e-4.

As depicted in Figure 1F, the N-terminal extensions of Vangl2-Long isoforms contain several residues that are strictly conserved in vertebrates. As a first step towards a functional characterization of these highly conserved residues, two of them, namely V8 and L11, were targeted by mutagenesis with the rationale that hydrophobic residues frequently contribute critical roles in the context of membrane proteins. Accordingly, V8 and L11 were substituted for alanines either individually (V8A or L11A) or simultaneously (V8A L11A) in GFP-tagged Vangl2-Long. IMCD3 cells stably transfected with either of these wild-type or mutant constructs were subsequently analyzed by IF microscopy. Careful examination and quantification of our IF data revealed a moderate impact of the V8A mutation on the cellular distribution of Vangl2-Long. In particular, recruitment of the V8A mutant protein at the plasma membrane was more conspicuous as compared to the wild type protein (Figure 6A first and second rows). Another feature of the V8A phenotype was the reduced colocalization of the mutant protein with GM130 positive membranes (Figure 6A and 6B). The single L11A and double V8A L11A mutations, on the other hand, led to similar localization defects but with a strikingly and significantly higher penetrance as judged by the dramatic accumulation of both mutant proteins at E-cadherin-containing cell-cell contacts and the concomitant decrease of the long isoform in the Golgi apparatus (Figure 6A, third and fourth rows and Figure 6B). These data further illustrate the key role played by the N-terminal extension of Vangl2-Long in promoting its targeting or retention to the Golgi apparatus. They also point to a differential contribution of two highly conserved residues to the regulatory mechanism involved.

**Figure 6.**
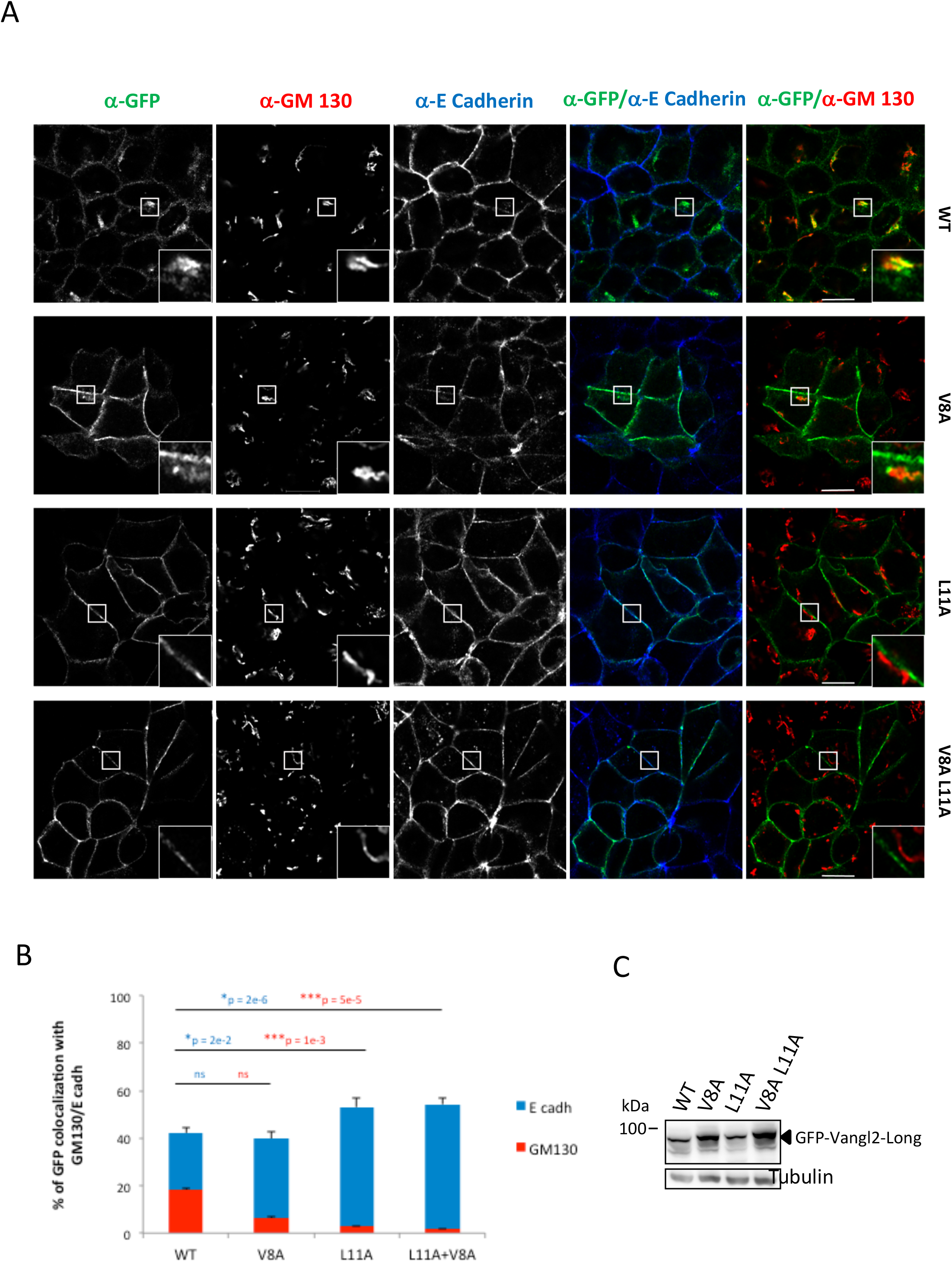
Val8 and Leu11 are two highly conserved residues of the N-terminal extension required for the Golgi association of Vangl2-Long. **(A)** IMCD3 cells were stably transfected with a set of GFP-Vangl2-Long constructs harboring either a wild-type V8 L11 sequence (first row, WT), a single V8A (second row, V8A) or L11A (third row, L11A) substitution or a double V8A, L11A (fourth row, V8A L11A) mutation. Cells were analyzed by immunofluorescence microscopy using antibodies directed against GFP, E-cadherin and GM130. Scale bars represent 10 μm **(B)** Quantification of the % of WT and the indicated mutant proteins colocalizing with E-cadherin (blue histograms) and GM130 (red histograms) using MATLAB code. Histograms represent the mean ± s.e.m calculated for three independent experiments (>30 cells each). P-values evaluated by Student’s t-test. ns p>0.05, * p<0.05. ** p<1e-3, *** p<le-4. All statistical tests are compared to WT. **(C)** Protein extracts from the same cells as used in (A) were probed by Western blotting with mAb 36E3 (upper panel) and anti-tubulin (lower panel) antibodies.

### Vangl2-Long expression in neural plate cells of *Xenopus* embryos

In the absence of a robust experimental PCP model in mammalian cultured cells, we turned to the early embryonic development in *Xenopus* to address the functional relevance of Vangl2-Long in the context of this signaling pathway. We first verified that mAb 36E3 could detect xVangl2 but not xVangl1 by Western Blot (Figure S9A) before assessing the existence of a frog Vangl2-Long isoform in kidney epithelial *Xenopus* A6 cells. This provided a first piece of evidence as mAb 36E3 detected, like in SKBR7 cells, two distinct bands of 62kDa and a 70kDa (Figure 7A). Furthermore, mass spectrometry analysis of *Xenopus* embryo proteins immunoprecipitated by the Vangl2 antibody recovered several peptides of the xVangl2A and/or Vangl2B isoforms (Figures 7B and table S1). Two peptides were of particular interest here as one spanned the expected Met-initiated N-terminus of xVangl2 while the other contained an additional stretch of four N-terminal aa whose sequence perfectly matched the last four residues of the predicted N-terminal extension of xVangl2-Long (Figures 7C and table S1).

**Figure 7.**
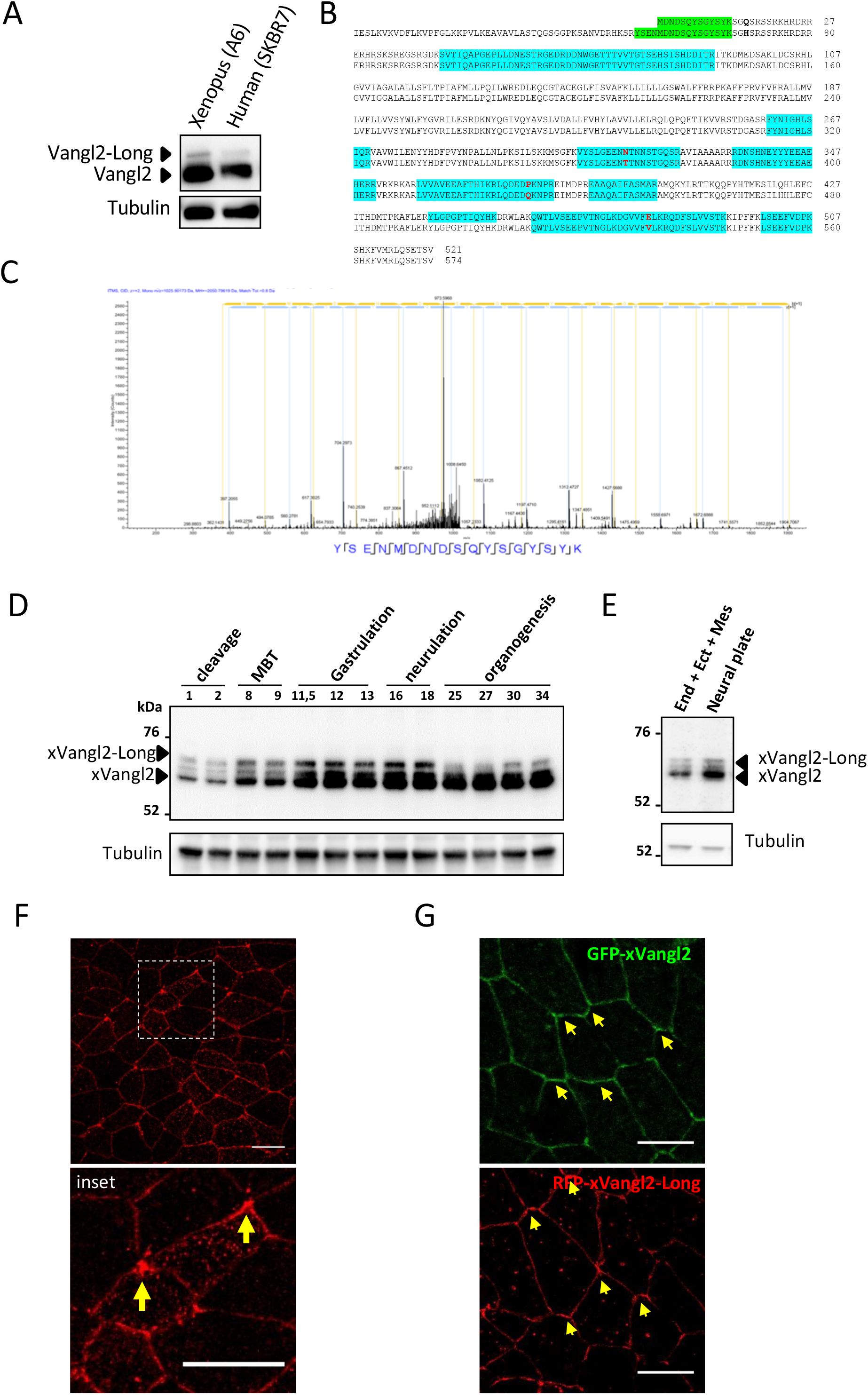
Biochemical and cytological characterization of ×Vangl2-Long during gastrulation and neurulation in *Xenopus* embryos. **(A)** Western blot analysis of *Xenopus* A6 and human SKBR7 cells using mAb 36E3. Immunoblotting with an anti-tubulin antibody is shown in the lower panel and can be used as a protein loading control. Note that in *Xenopus* A6 cells mAb 36E3 also recognizes two polypeptides, a major 62 kDa and a less intense 70 kDa band, that in SKBR7 correspond to Vangl2 and Vangl2-Long, respectively. **(B)** Peptide coverage of xVangl2 as determined by mass spectrometry analysis. Sequences (in blue) corresponding to both Vangl2A (upper) and 2B (lower) xenopus isoforms were detected by LC-MSMS. Slight differences in amino acids are indicated in red. Both MDNDSQYSGYSYK canonical Nterminus sequence or YSENMDNDSQYSGYSYK peptide that highlighted an N-terminus extension sequence were identified and are indicated in green. **(C)** ms/ms spectrum of the YSENMDNDSQYSGYSYK peptide MH+: 2050.7961 Da **(D)** Whole *Xenopus* embryos collected at the indicated stages of development were used for protein extraction and western blot analysis with mAb 36E3 (upper panel). Immunoblotting experiments with an anti-tubulin antibody (lower panel) were performed in parallel for protein loading control. MBT, Midblastula transition. **(E)** Embryos at stage 14 were dissected to isolate the neural plate (neural plate) from the rest of the embryo that comprised endodermal, ectodermal and mesodermal tissues (End + Ect + Mes). Proteins of both explants were extracted and probed by Western Blotting with mAb 36E3 (upper panel) and anti-tubulin (lower panel). The bands corresponding to xVangl2 and xVangl2-Long are indicated by black arrows. **(F)** Immunofluorescence microscopy analysis of *Xenopus* embryo neural plate cells stained with mAb 36E3. White dotted line square indicates the region magnified in the lower panel. Arrows indicate the polarized accumulation of endogenous xVangl2 at the anterior pole of each cell. **(F)** Embryos injected into two dorsal blastomeres at four-cell stage with GFP-xVangl2mRNA or RFP-xVangl2-Long mRNA as indicated. Arrows indicate the slightly polarized distribution of GFP-xVangl2 and RFP-xVangl2-Long along the antero-posterior axis in each cell. Scale bars= 20 μm.

The biochemical data shown above together with our bioinformatics analysis of the 5’-UTR region of the xVangl2 loci strongly argue in favor of an alternatively translated xVangl2-Long isoform in this species. It should be noted here that our mass spectrometry analysis of the polypeptides immunoprecipitated by mAb 36E3 also recovered four xVangl1 peptides, further extending the interaction data obtained in mouse and human cells (Table S2) (11). Immunofluorescence experiments done in IMCD3 cells, revealed that GFP-xVangl2 and GFP-xVangl2-Long exhibited a very similar localization pattern at the plasma membrane and in the Golgi compartment as observed with the human Vangl2 isoforms (Figure S9B). In summary, our data confirm the existence of xVangl2 and xVangl2-long in *Xenopus* embryos and illustrate the conserved localization properties of these proteins when expressed in murine epithelial cells.

In order to document the expression pattern of both xVangl2 isoforms during the early developmental stages of *Xenopus* embryos, we monitored their levels at different time points during development, from cleavage to neurulation (Figure 7D). This time-course study revealed that both isoforms are detectable from the beginning of embryogenesis (stage 1). At mid-blastula transition (MBT, stages 8-9), expression of both xVangl2 and xVangl2-Long starts to increase, reaching highest levels during gastrulation and neurulation. xVangl2 remained highly expressed from gastrulation to organogenesis. In contrast, a significant decline in xVangl2-Long levels occurred during stages 25-27 before rising at later stages of organogenesis (stages 30-34). These results therefore indicate that xVangl2 expression and the alternative translation of xVangl2-Long are both subjected to a precise developmentally regulated program. To gain knowledge about the tissue distribution of xVangl2-Long, we probed by immunoblotting extracts from explanted neural plates with our Vangl2 antibody (Figure 7E). xVangl2 and xVangl2-Long were found to be slightly enriched in the neuroectoderm relative to the surrounding tissues. Altogether, these results suggest that xVangl2-Long isoform could have a specific role during gastrulation and neurulation.

Next, we exploited the immunoreactivity of mAb 36E3 antibody towards the *Xenopus* Vangl2 proteins to perform IF experiments in order to monitor the distribution of the endogenous proteins in the neural plate isolated at stage 14. We observed the presence of endogenous xVangl2 at the plasma membrane with a strong accumulation of the signal at the anterior edge of the cells (Figures 7F), consistent with the planar polarized localization of this protein previously reported during neurulation (48). We also examined the fate of ectopically expressed GFP-xVangl2 and compared it to the behavior of RFP-xVangl2-Long at the same neurulation stage (Figures 7G). This experiment showed the localization of GFP-xVangl2 at the plasma membrane as well as its slight enrichment at anterior edge of the cells, as observed with endogenous xVangl2, albeit to a much lesser extent. In this context, RFP-xVangl2-Long did not show strong intracellular staining as previously documented in IMCD3 cells but was predominantly found at the plasma membrane and in a slightly asymmetric manner reinforcing the idea that xVangl2-Long may play a role in neurulation.

### xVangl2-Long is required for convergent extension movements and neural tube closure

To determine whether ×Vangl2-Long, like xVangl2, has a morphogenetic function during embryogenesis we implemented a loss-of-function assay with a set of independent morpholino oligonucleotides (MOs) each targeting a different region of xVangl2 (MO Vangl2) or xVangl2-Long (MO(1) Vangl2-Long and MO(2) Vangl2-Long in Figure 8). MOs specificity was assessed by Western Blot analysis with mAb 36E3 (Figure 8A and 8B). As expected MO Vangl2 reduced the levels of xVangl2 by more than 60%. It impacted, however, the expression of xVangl2-Long to some extents as well. In contrast, the activities of MO(1)- and MO(2) Vangl2-Long were more specific as judged by their high potency at down-regulating xVangl2-Long and by their minor impact on the expression of xVangl2 (Figures 8A and 8B).

**Figure 8.**
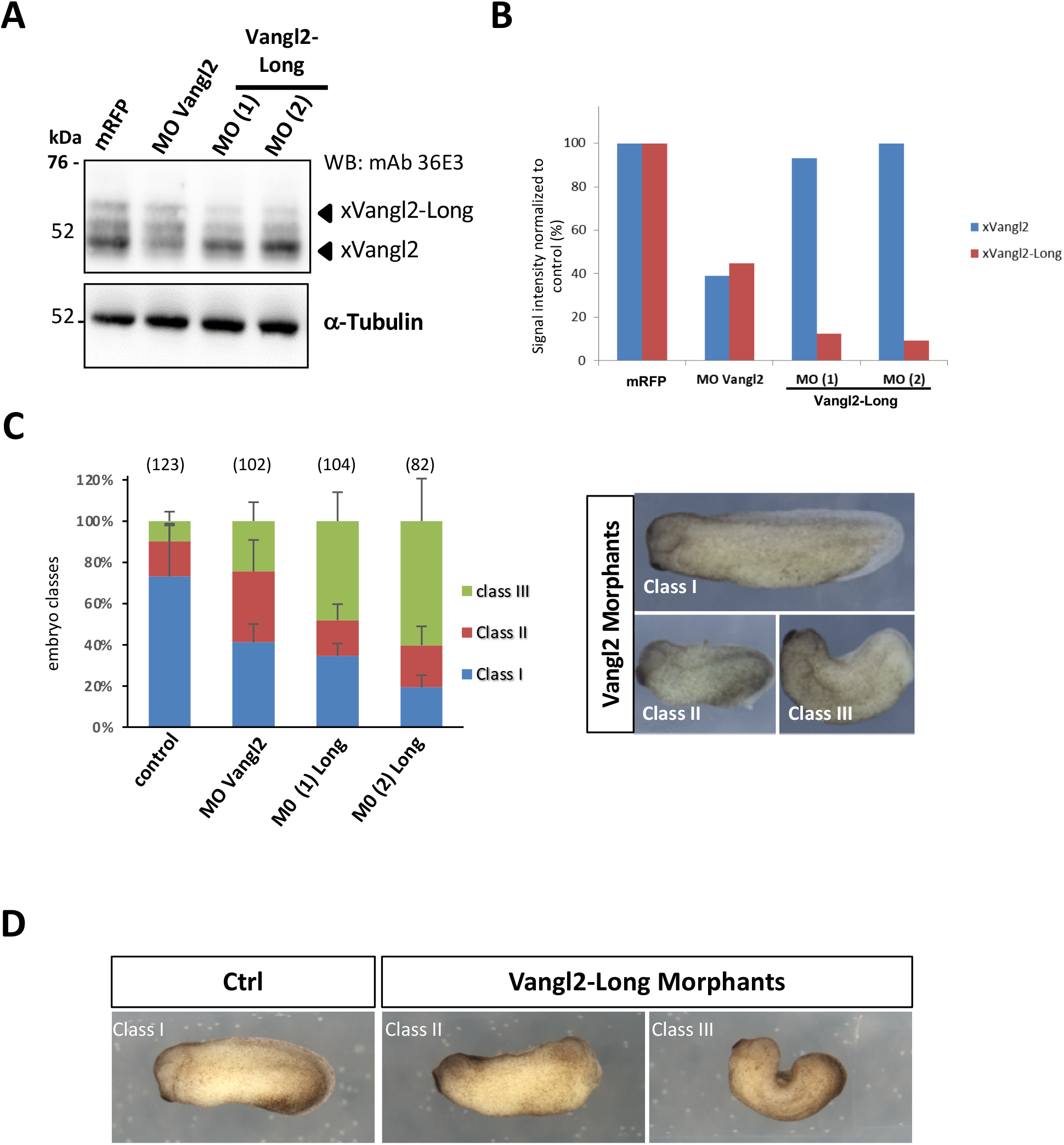
Vangl2-Long knockdown causes strong morphogenetic defects in Xenopus embryos. **(A)** *Xenopus* embryos were injected with mRFP, a morpholino-oligonucleotide (MO) directed against Vangl2 (MO Vangl2) or two independent MOs targeting Vangl2-Long (MO (1) and MO (2) Vangl2-Long) in dorsal blastomeres at 2-cell stage and grown until st. 13. Western blot analysis was carried out with mAb 36E3 (upper panel) and anti-tubulin (lower panel) antibodies. **(B)** Quantification of xVangl2 (blue) and xVangl2-Long (red) signals detected in (A). Note the efficient knockdown of xVangl2-Long achieved with the two independent MOs directed against this isoform while xVangl2 expression is largely unaffected. **(C)** Embryos were injected dorsally at 2-cell stage with either mRFP mRNA (control), MO Vangl2 or one of the two Vangl2-Long MOs as indicated. The histogram shows the class distribution among embryos injected with the indicated MO. Bars represent the proportions of embryos in each class (means ± s.e.m calculated for 5 independent experiments) as illustrated in the right panel with the phenotypes elicited by MO-Vangl2. Total numbers of embryos analyzed for each condition are given above each bar. **(D)** Morphologies of control *Xenopus* embryos injected with mRFP mRNA (Ctrl) or Vangl2-Long morphants injected with MO(1) Vangl2-Long (class II and class III).

It has been reported that xVangl2 is important for normal convergent extension movements and has a main role in the PCP pathway. In those works, the authors used MO Vangl2 and described that Vangl2 morphants displayed two different classes of defects (class II and class III) (18,19). The embryos with class II defects look shorter than controls. In the class III defects, the embryos are severely bent and do not close the neural tube (Figure 8C). We compared the effect of the MOs against xVangl2-Long with Vangl2 morphants (Figure 8C). We observed that the two MOs against Vangl2-Long yielded a high proportion of class II and class III embryos. Interestingly, Vangl2-Long morphants had more pronounced neural tube closure defects in comparison with Vangl2 morphants (Figures 8C and 8D). These results therefore strongly suggest that xVangl2-Long contributes an important function during embryo morphogenesis, particularly for normal neural tube closure.

## Discussion

Here, we report on a novel Vangl2 isoform, Vangl2-Long, that we identified in human, mouse and *Xenopus* and which bears an evolutionarily conserved N-terminal extension of 48aa in mammals and 53aa in frog. In agreement with a previous bioinformatics study (31), our data establish that translation of the human Vangl2-Long isoform requires a near-cognate in frame AUA alternative translation initiation site located 144 nucleotides upstream of the canonical *AUG* start codon of Vangl2. While detected in all Vangl2 positive cell lines tested so far, this 70kDa Vangl2-Long isoform is always expressed at lower levels than the canonical 62kDa Vangl2 protein. Interestingly, the expression pattern of Vangl2-Long in developing *Xenopus* embryos suggests that in a more physiological context, translation initiation of Vangl2-Long from the alternative AUA start codon is under the control of a developmentally regulated program. Our findings that isoform-specific MOs against Vangl2-Long impair convergent extension and neural tube closure is a strong indication that this new isoform contributes a critical function to the PCP signaling pathway that control these key morphogenetic events.

The remarkably high degree of conservation exhibited by the N-terminal extensions of all vertebrate members suggests that the capacity to encode a longer Vangl2 isoform has been acquired concomitantly with a gene duplication event that gave rise to the vertebrate *Vangl1* and *Vangl2* paralogs. Vertebrates have a nervous system totally different from invertebrates as most of the latter organisms, except hemichordate, urochordata and cephalochorda, do not have a neural tube. The neural tube becomes a physiologically sealed system from a very early developmental stage only in vertebrates (49). This suggests that the N-terminal extension of Vangl2 may have provided vertebrates with a novel morphogenetic property, a contention that is fully supported by the finding that Vangl2-Long is necessary for proper neural tube closure in *Xenopus* embryos.

Along the same line, mutations in the human *Vangl2* and *Vangl1* genes have been linked to genetic diseases characterized by neural tube closure defects (12,13). Given the strong PCP phenotypes elicited by loss-of-function of xVangl2-Long in *Xenopus* embryos, we wondered whether mutations associated with severe forms of neural tube closure defects or other human developmental disorders related to PCP defects could be mapped to the N-terminal extension of Vangl2-Long. However, a search for genetic variants in NTD patient cohorts failed to identify variants that specifically affect the coding sequence of the Vangl2-Long extension (E. Ross and V. Agular-Pulido, personal communications). A possible explanation is that human embryos with mutations in the extension of Vangl2-Long may not survive due to the functional importance of Vangl2-Long for neural tube closure. Testing this hypothesis will certainly deserve further investigations using larger cohorts of NTD patients as well as by performing a genetic analysis of mis-carried human embryos.

The capacity of Vangl1/2 to heterodimerize has been previously documented (11). Data presented here extend this biochemical property to the new Vangl2-Long isoform and reveal that the three proteins can form tripartite complexes in human cells possibly at their site of biosynthesis, i.e. the endoplasmic reticulum (Figure S10). Vangl1 and Vangl2 also form binary complexes in murine cells. Although the relative stoichiometry of Vangl1 within these complexes cannot be accurately determined, we could evaluate the molecular ratio between the two Vangl2 isoforms thanks to their common mAb 36E3 epitope. Thus by comparing the intensity of the 62kDa and 70 kDa signals we could estimate the Vangl2:Vangl2-Long ratio to range from 3:1 to 4:1. It follows that the minimal subunit composition of the Vangl1:Vangl2:Vangl2-Long tripartite complex should vary from 1:3:1 and 1:4:1, assuming the presence of only one molecule of Vangl1 in the multimeric complexes. This echoes the recently described supramolecular organization of fly core PCP complexes in which up to six Vang/Stbm molecules have been shown to form signalosome-like structures (50). Thus, while the molecular organization of Vang/Stbm and Vangl proteins appears to be widely conserved through evolution, the subunit composition has greatly evolved and increased in complexity with the appearance of multiple Vangl paralogs and isoforms.

Our localization data reveal that both isoforms can localize to the cell junctions of epithelial cells when expressed as GFP fusion proteins. Yet a significant fraction of GFP-Vangl2-LONG also localizes to the Golgi apparatus, suggesting that the N-terminal extension likely acts as a Golgi retention or localization signal. Several findings reported here indeed argue in favor of this possibility. First the finding that fusing the 48aa extension to the N-terminus of Vangl1 causes a dramatic redistribution of the chimeric GFP-Vangl1-Long protein from the plasma membrane to the Golgi apparatus, indicate that this region likely acts as an autonomous Golgi localization signal. A second argument is provided by the phenotypic characterization of mutations of two highly conserved hydrophobic residues of the 48aa extension. L11A substitution prevents the Golgi co-localization of Vangl2-Long and causes the mutant protein to relocate to cell-cell contacts in a manner very similar to the localization observed with the non-extended Vangl2 isoform. A final argument comes from the finding that *Xenopus* Vangl2 and Vangl2-Long proteins, partition similarly to the two human Vangl2 isoforms between the Golgi and plasma membrane compartments when expressed in a heterologous epithelial cell model (Figure S8). While all these data point to a highly conserved targeting function of the N-terminal extension of Vangl2-Long proteins, the exact molecular mechanism at work has still to be elucidated. In particular, it remains to determine whether the extensive association of Vangl2-Long with the Golgi apparatus stems from a slower transit through the different sub-compartments of this organelle (Figure S11, a), to a retention mechanism (Figure S11 b) and/or to an active retrieval mechanism that redirects TGN-emanating Vangl2-Long vesicles back to this compartment (Figure S11 c). Several approaches, including yeast two hybrid, peptide pull-down or differential interaction screens, have been attempted in the aim of identifying extension-specific protein interactors. However, all our efforts have remained unsuccessful so far. It will be therefore worthwhile addressing these mechanistic issues with alternative methods and by investigating the possible involvement of lipid-mediated interactions. It will be also important to examine whether the N-terminal extension affects the interaction of Vangl2-Long with the Arfrp1/AP-1 machinery (Figure S11 d) previously described for Vangl2 export from the TGN ((27).

Regardless of the molecular mechanism involved, the increased colocalization of GFP-Vangl2-Long with *cis-* and *trans*-Golgi markers may reflect a specialized regulatory function of GFP-Vangl2 at this location, either alone or in the tripartite complex that this isoform can form with Vangl1 and Vangl2. For instance, Vangl2-Long could regulate the export of Vangl1/2 or other core PCP proteins, such as Celsr1, out of the Golgi apparatus, thereby controlling their final destinations and potentially contributing to the establishment of PCP asymmetry. Consistent with this idea, it has been shown that core PCP proteins have an interdependent localization (23) and that asymmetric sorting regulates planar tissue patterning (51). Furthermore, it has been reported that members of the Flamingo protein family can promote the plasma membrane localization of Vangl2 in a mouse keratinocyte model (20). Investigating whether Vangl2-Long regulates trafficking events of other core PCP proteins, particularly at the level of the Golgi apparatus should warrant further analyses. Given the key role of this organelle in protein glycosylation (52) and even tough none of the Vangl proteins have been reported to be glycosylated so far, it remains also formally possible that Golgi trafficking of Vangl2-Long enables associated proteins to undergo a number of maturation steps mandatory for their functions or proper targeting to specific sites of the plasma membrane.

The route taken by Vangl2 between the ER and the TGN was poorly documented until now. The RUSH data reported here provide the first experimental evidence that this isoform takes a canonical route through the Golgi after its release from the ER. Importantly these RUSH data also clearly demonstrate that, upon release from the ER, Vangl2-Long exits the Golgi compartment with a marked kinetic delay relative to Vangl2. The Golgi localization of xVangl2-Long was also expected in *Xenopus* tissue. However we were surprised to find that, when ectopically expressed in embryonic neural tissue, xVangl2-Long, unlike its human counterpart was primarily detected at cell junctions (Figure 7) and not at the Golgi apparatus. This difference may stem from fundamentally different biological properties of cultured cells and tissues rendering the Golgi accumulation of the long isoform more difficult to appreciate in neural tissue. In support of this hypothesis, ectopic expression of GFP-xVangl2 and GFP-xVangl2-Long isoforms in IMCD3 cells yielded the same differential localization pattern as the former protein accumulated almost exclusively at the plasma membrane while the latter was detected both at the plasma membrane and in the Golgi (Figure S8).

Several proteins have been shown to use non-AUG alternative initiation sites to generate isoforms with distinct biochemical or cellular properties, providing putative mechanisms for further functional diversity (36). Vangl2-Long is no exception and its localization in the Golgi apparatus shows similarities to an N-terminally extended isoform of VEGF, called L-VEGF. L-VEGF has been reported to localize to the Golgi apparatus, whereas VEGF is secreted (44). The highly conserved Leu11 residue in the Vangl2 N-terminal extension is clearly important for Golgi retention (Figure 6). This observation is somewhat reminiscent of another human protein, NEFA, for which a Golgi retention signal located in its N-terminal Leu/Ile-rich region has been reported (53). As already mentioned above the N-terminal extension could also provide Vangl2-Long with the ability to interact with specific membrane lipids as recently described by Hopkins et al. for PTEN-Long, a secreted translational variant of PTEN that can translocate into neighboring tumor cells, where it induces tumor cell death (41), or to the mitochondria where it regulates mitochondrial energy metabolism (42).

## Acknowledgements

We thank Franck Perez and John Wallingford for the RUSH and *Xenopus* GFP-Vangl1/2 plasmids, respectively, Vita Bryja for the KO-Vang2 HEK 293T cell line, Frédérique Lembo for her help with plasmid construction, François Coulier for his help with bioinformatics analysis of the vertebrate N-terminal extensions, Olivier Rosnet for WB on A6 cells, Avais Daulat, Rania Ghossoub, Paula Michea, Michael Sebbagh and all members of the Jean-Paul Borg and Pascale Zimmerman teams for their stimulating discussions and helpful advices, members of the CRCM mass spectrometry and cytometry facilities, members of the MiMabs company for their contribution to the production of Vangl1/2 mAbs. This work was funded by La Ligue Nationale Contre le Cancer (Label Ligue JPB), Institut Universitaire de France and PL BIO Institut National du Cancer (grant INCa-9474). A.W. is a recipient of a fellowship from Assistance Publique des Hôpitaux de Marseille et Aix-Marseille Université. Proteomic analyses were done using the mass spectrometry facility of Marseille Proteomics (marseille-proteomique.univ-amu.fr) supported by IBISA (Infrastructures Biologie Santé et Agronomie), Plateforme Technologique Aix-Marseille, the Cancéropôle PACA, the Provence-Alpes-Côte d’Azur Région, the Institut Paoli-Calmettes and the Centre de Recherche en Cancérologie de Marseille, Fonds Européen de Developpement Regional and Plan Cancer. Jean-Paul Borg is a scholar of Institut Universitaire de France.

## Author contributions

S.A. and L.C. performed and processed mass spectometry analysis. D.I. and A.S. contributed to data processing of the IF and RUSH experiments. S.M., A.W. and E.B. generated anti-Vangl1 and anti-Vangl2 antibodies. T.M.P. contributed to the design of N-Vgl2 pAb and to scientific discussions. L.K. contributed to scientific discussions and supervised the *Xenopus* work performed by D.R., both wrote the corresponding part of the manuscript. A.W. contributed to all experiments. E.B. constructed expression plasmids, contributed to some biochemical experiments and supervised with JPB this study. A.W., E.B. and J.P.B wrote the manuscript.

## Material and Methods

### Cell culture, transfections, stable cell lines and generation of K.O cells lines using CRISPR/Cas9

Most cell lines used in this study were from and grown as recommended by American Type Culture Collection in the presence of 100Uml_l of penicillin, 100 mg ml_1 of streptomycin and 10% heat-inactivated fetal bovine serum. KO-Vangl2 HEK 293T cells were kindly provided by Vita Bryja (Institute of Experimental Biology, Faculty of Science, Masaryk University, Brno, Czech Republic). Murine epithelial IMCD3 cell line was cultured in DMEM/F12 growth medium, HEK 293T cells were propagated in DMEM medium and SKBR7 cells were grown in RPMI medium. Cell culture was performed at 37°C in 5% CO2 incubator. A6 cells were grown at 27°C in 55% Leibovitz’s L15 medium, 20 u/ml penicillin, 20 μg/ml streptomycin (Life technologies). For RNAi experiments, cells were transfected with 20nM RNAi using Lipofectamine RNAiMAX reagent (Life technologies); cells were analyzed 48 or 72 hours after RNAi treatment. We used polyethyleneimine (PEI) for transient transfection of DNA constructs in HEK 293T cells. For stable cell lines, IMCD3 cells were transfected with DNA constructs using Lipofectamine Plus reagent (Life technologies) according to manufacturer’s recommendations. Stable transfectants were selected in medium containing 1 mg/ml G418 (Life technologies) for 2 weeks. Crispr-Cas9 mediated gene editing of Vangl2 locus in IMCD3 cells was achieved by cloning a guide RNA (gRNA) designed by Life technologies to target the second exon of murine Vangl2 (gRNA: 5’-TCGGCTATTCCTACAAGTC-3’) into pSpCas9(BB)-2A-GFP (PX458, Addgene #48138). Upon transfection, GFP-positive IMCD-3 cells were sorted and individually seeded in 96 well plates by Fluorescence activated cell sorting (FACS) using an ARIA III (Becton Dickinson) cell sorter. Clones were then expanded and screened by Western blot using mAb 36E3.

### Antibodies

Rat anti-Vangl2 antibodies (mAb 2G4 and mAb 36E3) were obtained as described elsewhere (11). N-Vgl2 pAb was obtained by immunizing two rabbits with an 18mer peptide sequence (PKRPQPAALERYKARRSD) spanning aa30-47 of the N-terminal extension of Vangl2-Long. Immunoglobulins that specifically react against Vangl2-Long were obtained by affinity purification of N-Vgl2 pAb on beads covalently coupled to the immunogenic peptide describesd above. Mouse anti-α-Tubulin (T9026) was from Sigma. Rabbit (A11122) and mouse (JL-8) anti-GFP antibodies were from Life technologies and Takarabio respectively. Mouse anti-GM130 (610822) was from BD Transduction Laboratories. Rabbit anti-KDAEL (ab176333) was from Abcam. Goat anti-E-cadherin (AF648) was from R & D Systems. Secondary antibodies were from Life technologies.

Mouse anti-Vangl1 monoclonal antibodies were produced in collaboration with Mimabs (Luminy - Marseille, France) by immunizing mice with a recombinant GST-tagged Vangl1 fragment spanning amino-acids 1-102. Hybridoma supernatants were counterscreened by ELISA against GST and GST-Vangl2. Clones that reacted specifically against GST-Vangl1 were selected for further biochemical and cytological characterization.

### Expression vectors and RNAi sequences

#### All expression plasmids used in this work are listed in table S3. Detailed informations about their production can be obtrained upon request

All RNAi targeting human sequences (siGENOME) and the non-targeting control RNAi (siCNT) were purchased from Dharmacon Inc. SiVGL1#1 (5’GAACAUGAACGGCGAGUAA3’), SiVGL1#2 (5’GGAAAUGAUUCUACUCGGA3’).

#### Immunofluorescence staining and confocal microscopy analysis

Cells grown on Rat collagen-coated glass coverslips were fixed for 10 min with 4% PFA in PBS, rinsed in PBS, permeabilized for 10 min with 0.4% Triton in PBS and quenched with 3% BSA in PBS. Cells were then incubated with the indicated primary antibodies overnight, washed and stained with secondary antibodies coupled to appropriate fluorochromes. Coverslips were mounted in ProlongGold antifade reagent with DAPI mounting media (Life technologies). Cells were then visualized and images acquired with a Carl Zeiss LSM880 confocal microscope using a Plan-Apochromat 63X oil immersion objective. Confocal image analyses were processed using Zeiss LSM Image browser version ZEN and Adobe Photoshop softwares.

#### Image analysis method for Vangl2/Vangl2-Long colocalization

Colocalization of Vangl2 with the Golgi marker GM130 as well as the membrane marker E-cadherin, was automatically computed with Matlab (The Mathworks). Confocal acquisitions of fixed cells were performed with an LSM880 microscope (Zeiss), with a calibration of 70 nm/pixel and 370 nm/z-plane. Custom Matlab scripts were developed for quantification, as follow: (i) z-stacks from green (Vangl2), red (GM130) and Magenta (E-cadherin) channels were first smoothed by a median filter of 3-pixel width. (ii) The resulting images were automatically thresholded using the function “opthr”, written by F.T. Marti, which can be downloaded from the Matlab file exchange website at mathworks.com/matlabcentral/fileexchange/132-opthr. (iii) Binary structures smaller than 100 voxels were discarded as non-specific. The resulting binary z-stacks allowed computing the percentage of Vangl2 colocalized with GM130 or E-cadherin staining as follow: percent of voxels segmented both for green (Vangl2) and red (GM130) or magenta (E-cadherin) channels relative to pixels segmented for the green channel. Thresholding procedures were validated by visual inspection. Quantification was performed on all voxels of the acquired confocal z-stack, allowing to properly take into account the 3D geometry of the Golgi apparatus.

The source code for image analysis is available as open-source software for academic and non-profit research upon reasonable request to A.S.

### Statistics

Each data set was first tested for normality using a one-sample Kolmogorov-Smirnov test. Since both data sets had at least 7 values and were both evaluated as normal, comparison was evaluated by a two-tail Student’s t-test, assuming equal variance. All statistics were performed with Matlab Statistical Toolbox (The Mathworks).

### RUSH experiments and live cell imaging

IMCD3 cells were plated in imaging chambers (35mm μ-dish, Ibidi) at a confluence of 50% and transiently transfected with GFP-Vangl2 and GFP-Vangl2 RUSH reporter contructs. Neutravidin (Life technologies) was added to culture media (100 nM in optiMEM and 40 nM in DMEM/F12) to prevent uncontrolled release of the cargo from the ER in the absence of exogenously added biotin. Cells were transduced 24 hours later with CellLight® Golgi-RFP (C10593, Life technologies) for 20 hours. Synchronous release of the GFP-tagged Vangl2 cargoes from the ER was induced by addition of 80 μM biotin to the medium and their traffiking was monitored by live cell imaging. Live cells were maintained at 37°C with 5% CO2 using a Pecon/Zeiss unit temperature and CO2 controller modules and a heating insert adapted to Zeiss Axio Observer Z1 microscope equipped with a CSU-X1-A3 Yokogawa spinning disk with dual camera port. Images were captured using an alpha Plan-fluar 100x (NA 1.45) oil objective. The laser rank was controlled using an iLas2 Roper Scientific Module (Roper Scientific SAS, Evry, France) driven by MetaMorph Software 7.8.13 (Molecular Devices, Berkshire, UK). During live cell imaging the two fluorescent channel images were acquired simultaneously on the two EMCCD Cameras Evolve 512 (Photometrics). For each time point position we acquired a Z-stack of 9 images separated by 0.8 mm and before biotin addition, we made an image defined as the initial image, T0. For colocalization evaluation the alignment of the two cameras was adjusted firstly mechanically on the Yokogawa spinning disk module using a control slide with multifluorescent beads, diameter of 3.0 mm (Rainbow Fluorescent Particle Slide from Spherotech). A second fine adjustment of the alignment was done at the beginning of the image processing.

Image processing and analysis were done with ImageJ software (National Institutes of Health, Bethesda, MD, USA). We wrote a macro to automatize the image processing using different plug-ins and parameter values to perform segmentation of the Golgi with as much precision as possible. First the two channel images were cropped to analyze cells one by one and to eliminate artefacts. Next, x,y positions of the two channels were adjusted in accordance with the control image using sherotech bead slide. We used medium filter plugin to reduce noise and used “find maximum plug-in” with 5 different values of “tolerance” parameter to perform optimal segmentation according to the signal to noise ratio, to quantify the colocalization between VangL2/Vangl2-Long (green channel) and the Golgi marker (red channel). We measured the average amount of green signal in the segmented image of Golgi, over the entire z-stack image for each time point. A normalization was applied to be able to compare the different conditions, using the following formula : A(t_j_)-A(t_o_)/A_max_(t_o_)-A(t_o_) where A(t_j_) is the average of green at a given time-point “i” (t_i_), A(t_o_) the average of green at initial time point t_o_, before biotin addition, and A_max_(t_x_) the maximum average amount of green (max colocalization) at time-point “x”. Images and videos were generated by maximum-projection of the z-stacks. Images and videos were generated by maximum projection of the z-stacks.

### Western blot (WB) analysis and IP assay

For Western blot analyses, cells were washed twice with ice-cold PBS and scraped immediately into ice-cold lysis buffer: 50mM Hepes-NaOH (pH=8), 150mM NaCl, 10% glycerol, 2mM EDTA and 0,5% NP-40 supplemented with a cocktail of protease inhibitors (Sigma-Aldrich). Cells were lysed during 15min at 4°C on a rotating-wheel and centrifugated at 13000rpm during 30 minutes. Protein concentration was determined using a Bradford assay. Cell lysates were mixed with 4X Laemmli sample buffer (in which βmercaptoethanol was added) in a 3:1 volume ratio, and boiled for 5 min. Proteins samples were loaded on NuPAGE™ 4-12 % Bis-Tris Gel (Life technologies) and separated by electrophoresis. Proteins were electrotransferred onto nitrocellulose blotting membranes (GE healthcare) and stained with Ponceau Red (Sigma Aldrich). The membranes were blocked with Tris Buffered Saline (TBS), 0.1% Tween 20, 5% (w/v) dried milk, incubated overnight with primary antibodies in blocking solution. Following extensive washing with TBS / 0.1% tween 20, the blots were incubated for 1 hour with the appropriate secondary HRP-conjugated antibody (Life technologies). HRP mediated chemiluminescence was detected using an ECL reagent kits (GE Healthcare) and signals were quantified by densitometry, using ImageJ.

Immunoprecipitation of GFP fusion proteins was done 48h post-transfection. 25-30uL of agarose beads covalently coupled to a GFP nanobody were prewashed with lysis buffer twice and incubated with 1-2 mg of lysates overnight at 4°C on a rotating-wheel. After extensive washing in lysis buffer, the bound material was eluted by the addition of Laemmli sample buffer 2X (in which βmercaptoethanol was added) and boiled for 5 min at 95°c. For immunoprecipitation of endogenous Vangl1, Vangl2 and Vangl2-Long, cell lysates were incubated overnight with indicated antibodies at 4°C on a rotating wheel. The immune complexes were then precipitated with either protein A agarose beads (GE Healthcare) or G for 4h at 4°C, washed 5 times in lysis buffer and analyzed by Western blot with the indicated antibodies.

### *Xenopus* embryo injections, plasmids, RNAs, and Mos

Eggs obtained from NASCO females were fertilized in vitro, dejellied and cultured as described previously (Marchal et al., 2009). Wild-type embryos were obtained using standard methods (Franco et al., 1999) from adult animals and staged according to Nieuwkoop and Faber (1994). Synthetic capped mRFP mRNA was produced using Ambion mMESSAGE mMACHINE Kit. pCS2-mRFP was linearized with NotI and mRNA was synthesized with Sp6 polymerase. 0,25ng of mRFP capped mRNA was used as injection control and tracer. GFP-xVangl1, GFP-xVangl2 and RFP-Vangl2 plasmids were kindly provided by J. Wallingford. 40 pg each of GFP-xVangl2 and RFP-Vangl2 capped mRNA were injected. Morpholino antisense oligonucleotides (MO) were obtained from Genetools®, and the sequences were as follows: MO Vangl2 5’-ACTGGGAATCGTTGTCCATGTTTC-3’; MO(1) Vangl2-Long 5’-CAACTTTAACCTTTAGCGACTCTAT-3’; MO(2) Vangl2-Long 5’-TAGCGACTCTATTTTGATTGGCTGT-3’. Embryos at 2-cell stage were injected dorsally with the following doses of MOs (MO Vangl2=30ng; MO(1) Vangl2-Long=40ng; MO(2) Vangl2-Long=40ng). Embryos were cultured in modified Barth’s solution until stage 14, when they were photographed and processed to observe at the microscope.

### *Xenopus* embryo microscopy

Confocal: Flat-mounted epidermal explants were examined with a Zeiss LSM 880 confocal microscope. Four-color confocal z-series images were acquired using sequential laser excitation, converted into single plane projection and analyzed using ImageJ software.

### Mass spectrometry analysis

Immunoprecipitated proteins were loaded on NuPAGE 4-12% Bis-Tris acrylamide gels (Life Technologies) to stack proteins in a single band that was stained with Imperial Blue (Pierce, Rockford, IL) and cut from the gel. Gels pieces were submitted to an in-gel trypsin digestion. Briefly, gel pieces were washed and destained using 100 mM NH4HCO3/acetonitrile (50/50). Destained gel pieces were shrunk with acetonitrile and were re-swollen in the presence of 100 mM ammonium bicarbonate in 50% acetonitrile and dried at room temperature. Protein bands were then rehydrated and cysteines were reduced using 10 mM DTT in 100 mM ammonium bicarbonate pH 8.0 for 45 min at 56 C before alkylation in the presence of 55 mM iodoacetamide in 100 mM ammonium bicarbonate pH 8.0 for 30 min at room temperature in the dark. Proteins were then washed twice in 100 mM ammonium bicarbonate and finally shrunk by incubation for 5 min with 100 mM ammonium bicarbonate in 50% acetonitrile. The resulting alkylated gel pieces were dried at room temperature. The dried gel pieces were re-swollen by incubation in 100 mM ammonium bicarbonate pH 8.0 supplemented with trypsin (12.5 ng/μL; Promega) for 1 h at 4 C and then incubated overnight at 37°C. Peptides were harvested by collecting the initial digestion solution and carrying out two extractions; first in 5% formic acid and then in 5% formic acid in 60% acetonitrile. Pooled extracts were dried down in a centrifugal vacuum system. Samples were reconstituted with 0.1% trifluoroacetic acid in 4% acetonitrile and analyzed by liquid chromatography (LC)-tandem mass spectrometry (MS/MS) using an Orbitrap Fusion Lumos Tribrid Mass Spectrometer (Thermo Electron, Bremen, Germany) both online with a nanoRSLC Ultimate 3000 chromatography system (Dionex, Sunnyvale, CA). Peptides were separated on a Dionex Acclaim PepMap RSLC C18 column. For peptide ionization in the EASY-Spray nanosource in front of the Orbitrap Fusion Lumos Tribrid Mass Spectrometer, spray voltage was set at 2.2 kV and the capillary temperature at 275 °C. The Orbitrap Lumos was used in data dependent mode to switch consistently between MS and MS/MS. Time between Masters Scans was set to 3 seconds. MS spectra were acquired with the Orbitrap in the range of m/z 400-1600 at a FWHM resolution of 120 000 measured at 400 m/z. AGC target was set at 4.0e5 with a 50 ms Maximum Injection Time. For internal mass calibration the 445.120025 ions was used as lock mass. The more abundant precursor ions were selected and collision-induced dissociation fragmentation was performed in the ion trap to have maximum sensitivity and yield a maximum amount of MS/MS data. Number of precursor ions was automatically defined along run in 3s windows using the “Inject Ions for All Available parallelizable time option” with a maximum injection time of 300 ms. The signal threshold for an MS/MS event was set to 5000 counts. Charge state screening was enabled to exclude precursors with 0 and 1 charge states. Dynamic exclusion was enabled with a repeat count of 1 and a duration of 60 s.

### Data Processing Protocol

Raw files generated from mass spectrometry analysis were processed with Proteome Discoverer 1.4 .1.14 (Thermo fisher Scientific) to search against the proteome reference of the *Xenopus laevis* protein database (56,145 entries, extracted on February 2018). The original fasta file was implemented with the long Vangl2 isoform by adding to the Nterminus part of the vangl2A sequence (accession number: Q90X64) the following 53 amino acids IESLKVKVDFLKVPFGLKKPVLKEAVAVLASTQGSGGPKSANVDRHKSRYSEN-. Database search with SequestHT were done using the following settings: a maximum of two trypsin miss cleavage allowed, methionine oxidation and N terminal protein acetylation as variable modifications, and cysteine carbamidomethylation as fixed modification. A peptide mass tolerance of 6 ppm and a fragment mass tolerance of 0.8 Da were allowed for search analysis. Only peptides with high Sequest scores were selected for protein identification. False discovery rate was set to 1% for protein identification.

---

**Figure S1.**
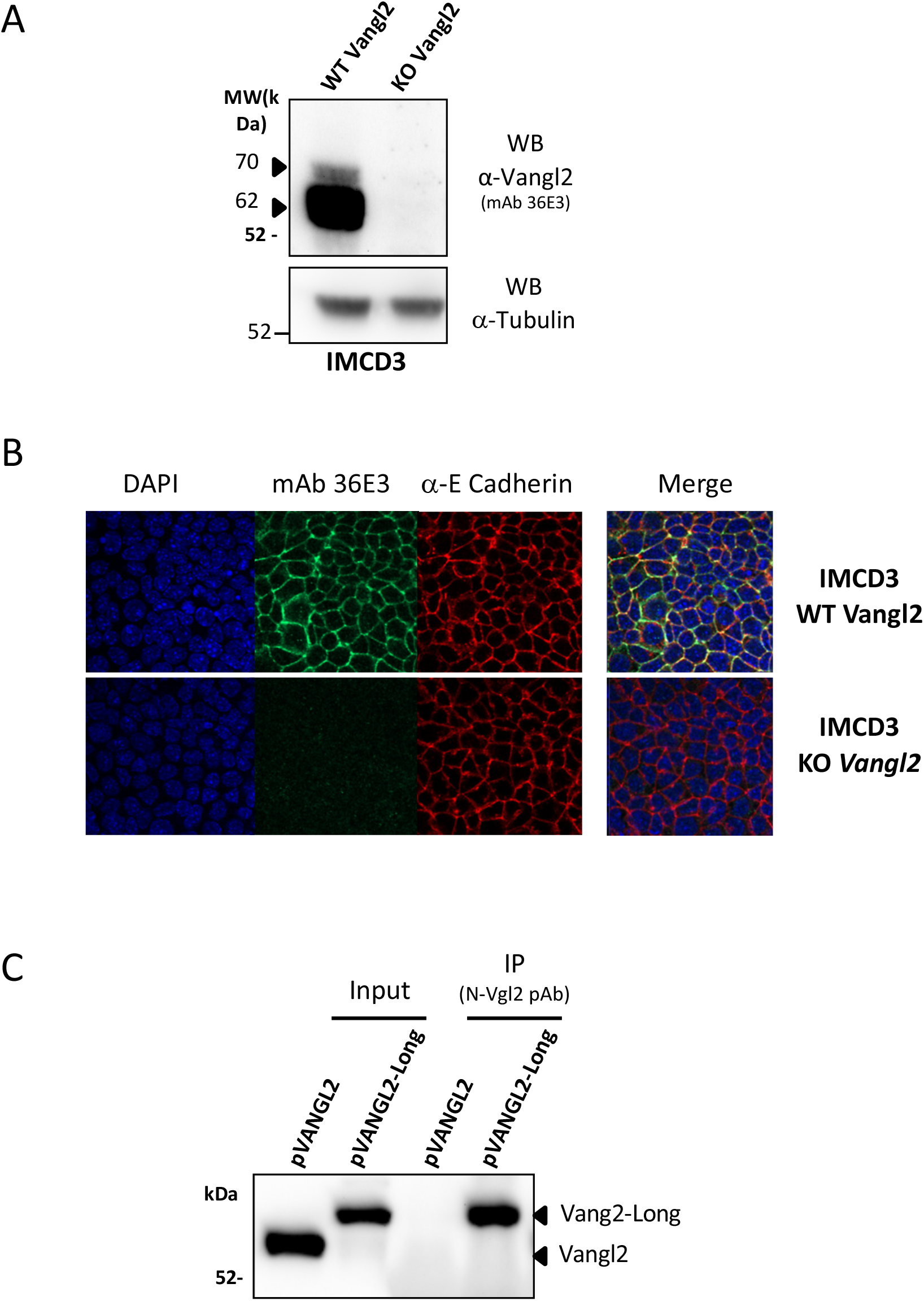
Validation of anti-Vangl2 (mAb 36E3) and anti-Vangl2-Long (N-Vgl2 pAb) antibodies. **(A)** IMCD3 cells with either a wild type (WT Vangl2) or Crispr-Cas9 generated knockout (KO Vangl2) Vangl2 alleles were assessed for Vangl2 expression by Western blot analysis with mAb 36E3. Note that Crispr-Cas9 mediated knockout of Vangl2 abrogates the expression of both the 62 kDa and 70 kDa proteins detected by mAb 36E3 **(B)** Immunofluorescence and confocal microscopy analysis of WT and KO Vangl2 IMCD3 cells were performed using the indicated antibodies. As expected, the IF staining observed with mAb 36E3 in control wild type IMCD3 cells is no longer detectable in KO Vangl2 mutant cells **(C)** KO Vangl2 IMCD3 cells stably transfected with pVangl2 or pVangl2-long were subjected to immunoprecipitation with N-Vgl2 pAb before Western blot analysis with mAb 36E3. Note the ability of N-Vgl2 pAb to immunoprecipitate Vangl2-Long but not Vangl2.

**Figure S2.**
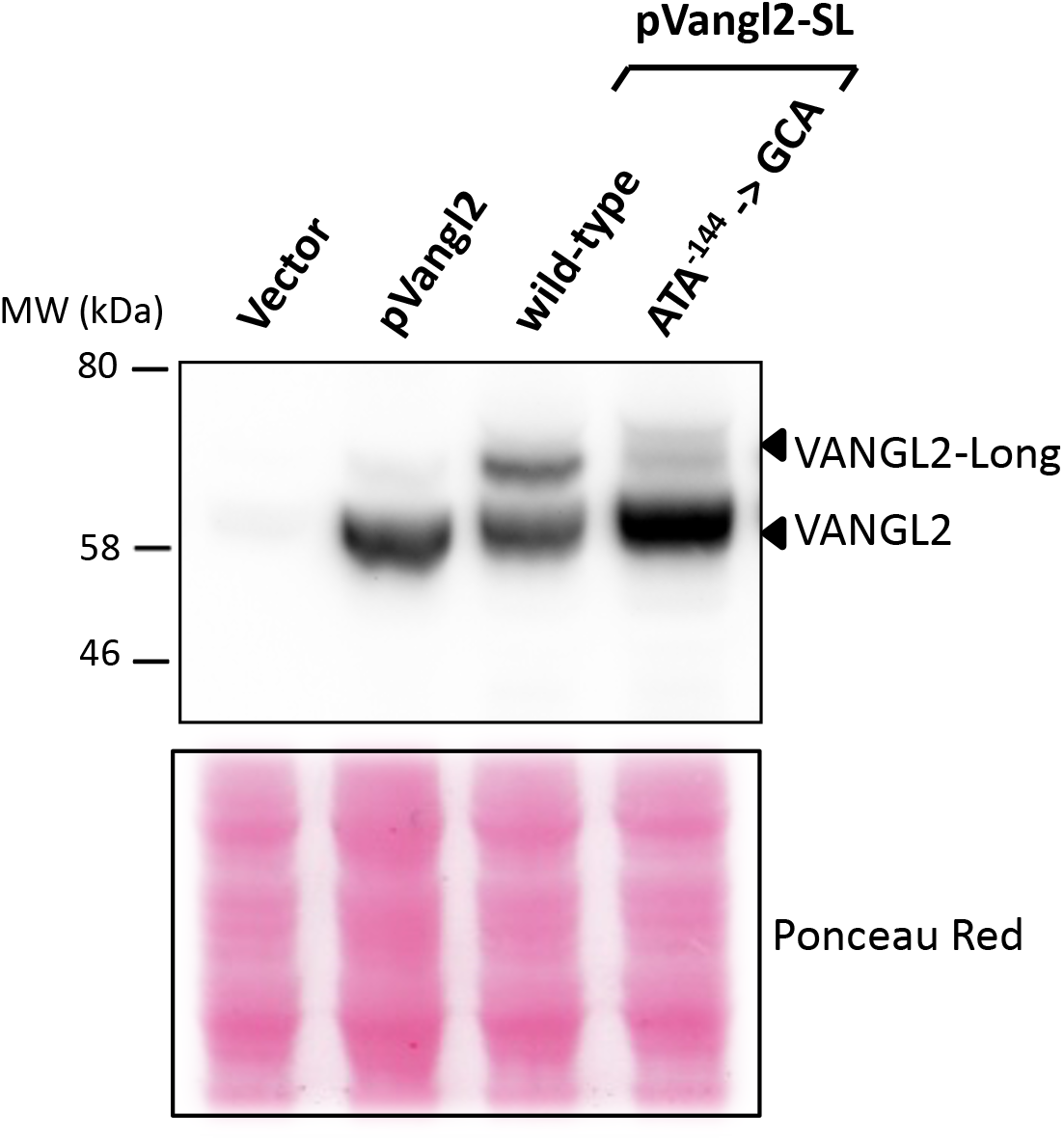
Vangl2-Long expression requires a near-cognate ATA alternative initiation site in position - 144. HEK 293T cells were transfected with either an empty plasmid (vector), pVangl2 or a pair of pVangl2-SL plasmids in which the putative non-conventional ATA’^144^ start codon was either left intact (wild-type) or mutated to GCA (ATA->GCA). Western blot analysis was carried out with mAb 36E3 to detect all Vangl2 isoforms. An image of the transfer membrane stained with Ponceau red is shown to confirm equal loading of the different samples.

**Figure S3.**
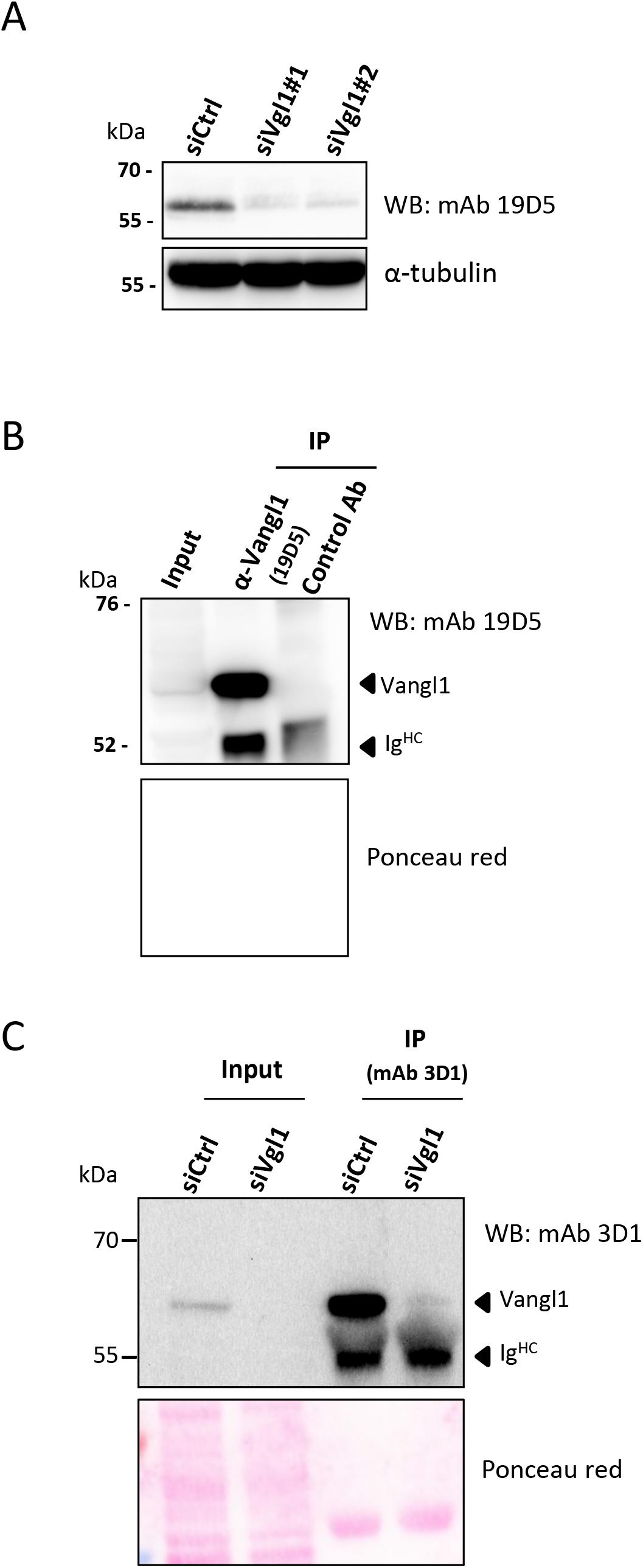
Validation of two Vangl1 specific monoclonal antibodies, mAb 19D5 and 3D1. **(A)** MCF7 cells were treated with two independent siRNA directed against human Vangl1 (siVgl1#1 and siVgl1#2) or a non-targeting siRNA (siCtrl). Total cell lysates were then analyzed by Western blotting with a novel Vangl1 specific monoclonal antibody, mAb 19D5, or with an anti-α tubulin used as a loading control **(B)** Immunoprecipitation of MCF-7 cell extracts with mAb 19D5 or a mouse isotypic control antibody (Control Ab). Protein IPs were analyzed by WB with the same mAb19D5 antibody. The lower panel shows an image of the transfer membrane stained with Ponceau red. **(C)** Murine epithelial (IMCD3) cells treated with Vangl1 siRNA (siVgl1) or a non-targeting siRNA (siCtrl) were processed for immunoprecipitation using a novel Vangl1 monoclonal antibody, mAb 3D1. Immunoprecipitated proteins were probed by WB with the same antibody. Arrows indicate the presence of the endogenous murine Vangl1 protein and the immunoglobulin heavy chain that is recognized by the secondary anti-mouse antibody used for the WB analysis. The lower panel shows an image of the transfer membrane stained with Ponceau red in order to confirm equal loading of each control and experimental samples.

**Figure S4.**
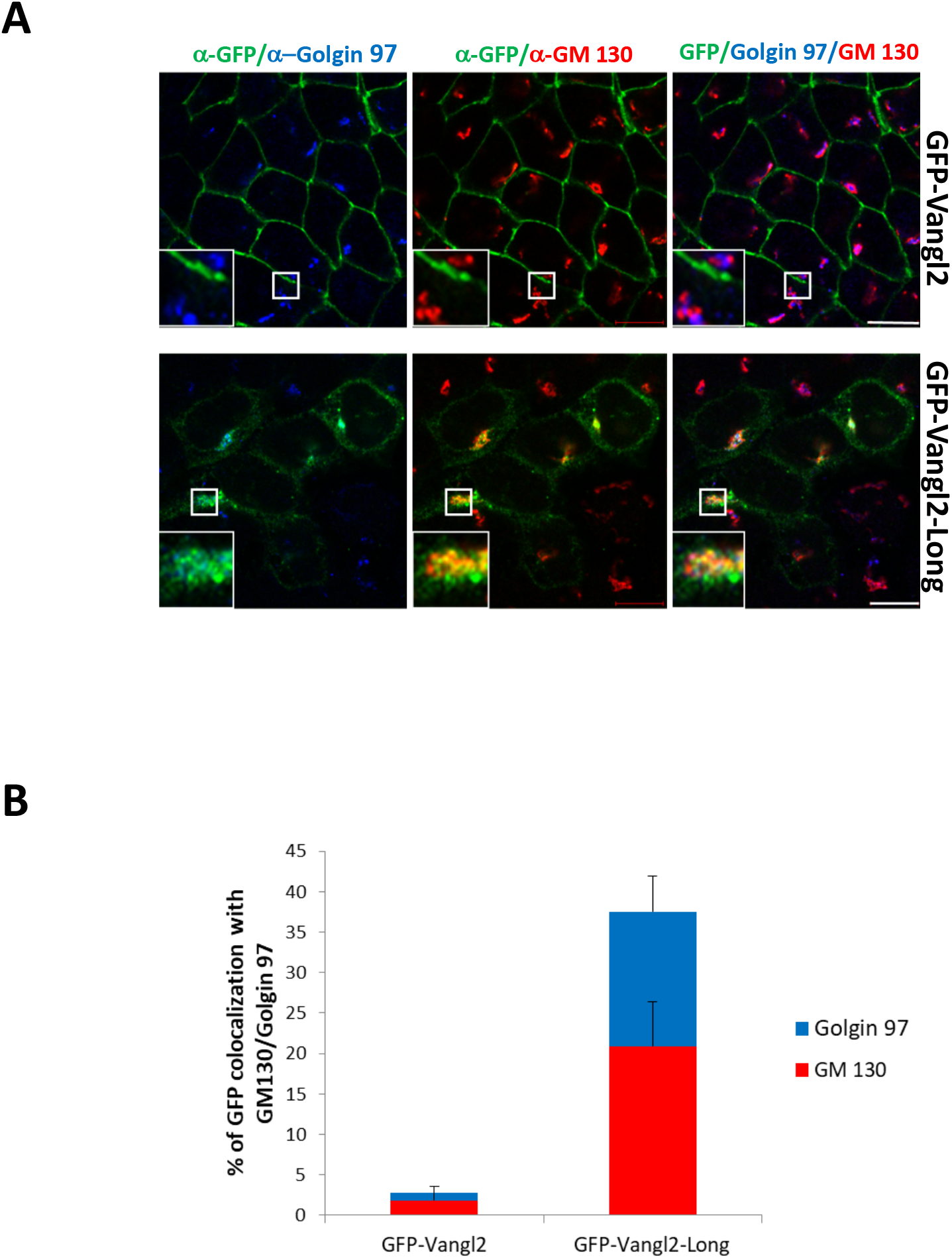
Partial colocalization of GFP-Vangl2-Long with the cis-Golgi GM 130 and trans-Golgi Golgin 97 markers. IMCD3 cells stably transfected with pGFP-Vangl2 or pGFP-Vangl2-Long were processed for immunofluorescence and confocal microscopy using the indicated antibodies Scale bar represents 10 μm. **(B)** Quantification of the % of GFP-Vangl2 and GFP-Vangl2-Long co-localizing with the cis-Golgi (GM130, red) and trans-Golgi (Golgin 97, blue) markers using MATLAB code. Histograms represent the mean ± s.e.m calculated for three independent experiments.

**Figure S5.**
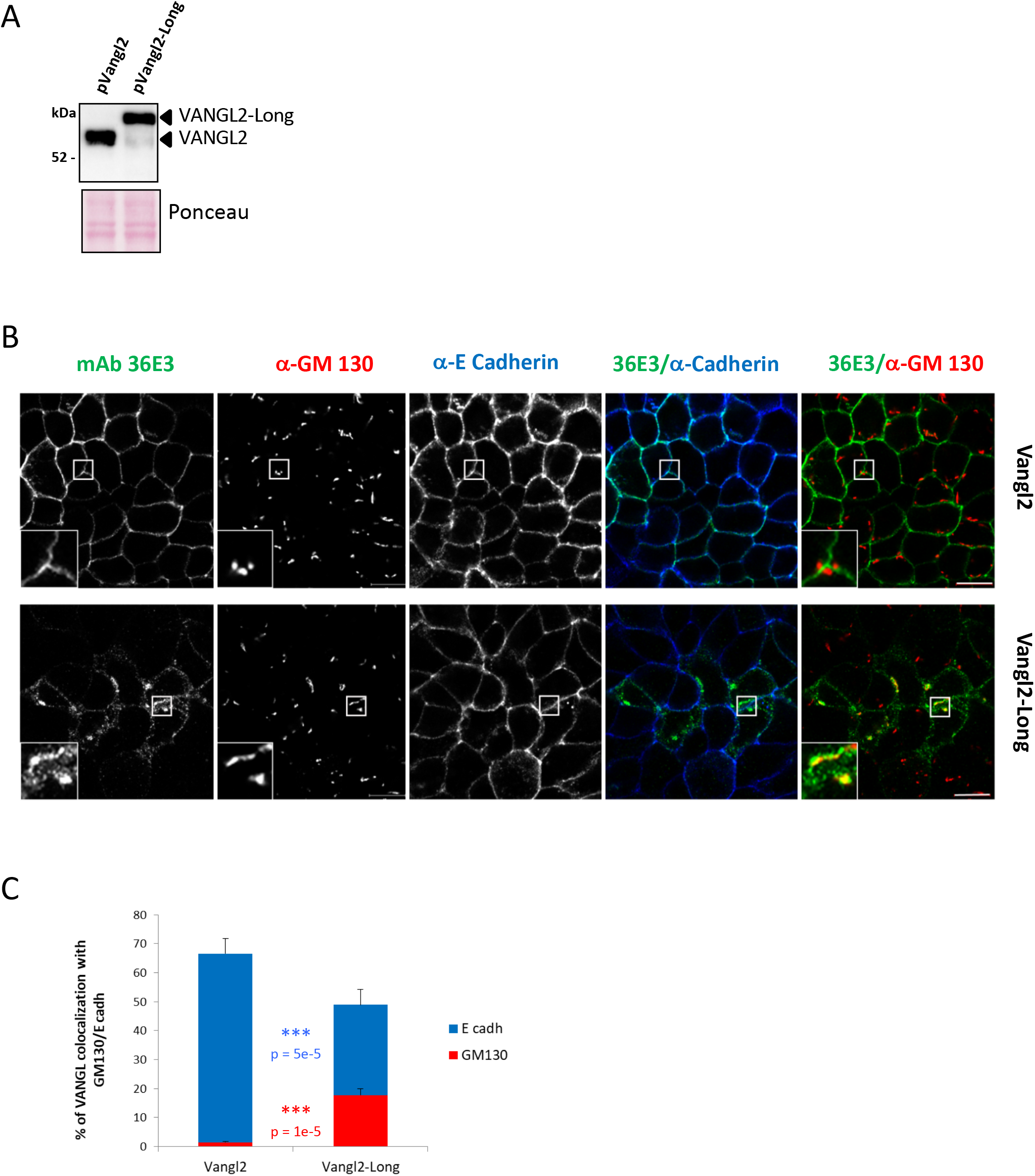
Localization of untagged Vangl2 and Vangl2-long in KO-Vangl2 IMCD3 cells. **(A)** KO-Vangl2 IMCD3 cells were stably transfected with the pVANGL2 or pVANGL2-Long plasmids described in Fig. 1D. Cell lysates were probed by Western blotting with the anti-Vangl2 mAb 36E3. Arrows indicate the positions of the relevant ectopically expressed Vangl2 isoforms. The lower panel shows an image of the transfer membrane stained with Ponceau red in order to confirm equal loading of each control and experimental samples. **(B)** Immunofluorescence and confocal microscopy analysis of cells described in (A) using the indicated antibodies. Note the extensive localization of Vangl2 at the plasma membrane and the substantial accumulation of untagged Vangl2-Long in the cis compartment of the Golgi apparatus decorated by GM130. Scale bars represent 10 μm. **(C)** Quantification of the % of VANGL2 and VANGL2-Long co-localizing with E-Cadherin (blue) and GM130 (red) fluorescence signals using MATLAB code. Histograms represent the mean ± s.e.m calculated for three independent experiments. P-values evaluated by Student’s t-test. *** p<1e-4.

**Figure S6.**
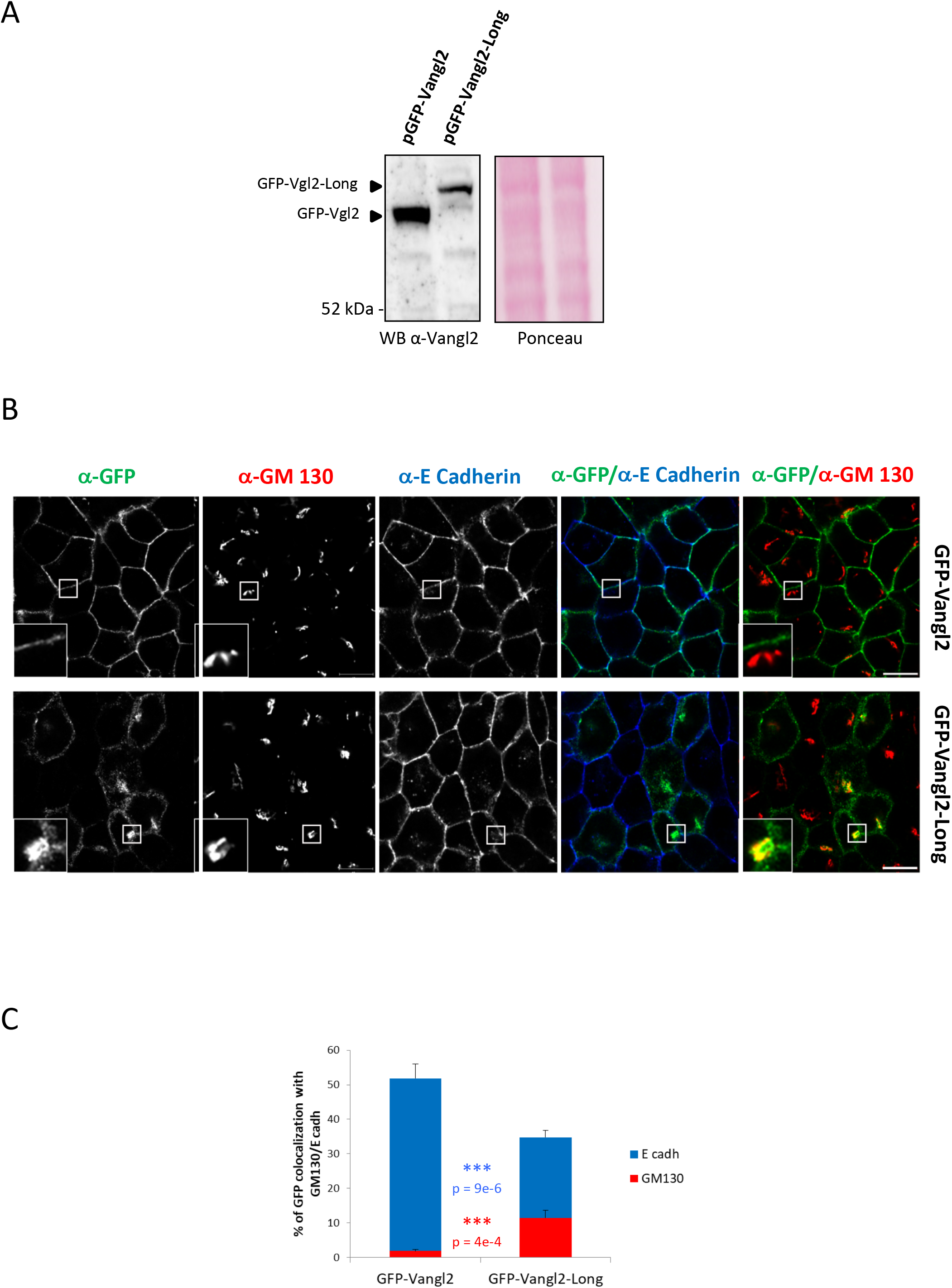
Golgi localization of Vangl2-Long in KO-Vangl2 IMCD3 cells. **(A)** Protein extracts from KO-Vangl2 IMCD3 cells stably transfected with pGFP-Vangl2 or pGFP-Vangl2-Long were probed by Western blotting with the anti-Vangl2 mAb 36E3. Arrows indicate the positions of GFP-Vangl2 and GFP-Vangl2-Long. **(B)** Immunofluorescence and confocal microscopy analysis of cells described in (A) using the indicated antibodies. Scale bars represent 10 μm. **(C)** Quantification of the % of GFP-VANGL2 and GFP-VANGL2-Long co-localizing with E-Cadherin (blue) and GM130 (red) fluorescence signals using MATLAB code. Histograms represent the mean ± s.e.m calculated for three independent experiments (>30 cells each). P-values evaluated by Student’s t-test. *** p<1e-4.

**Figure S7.**
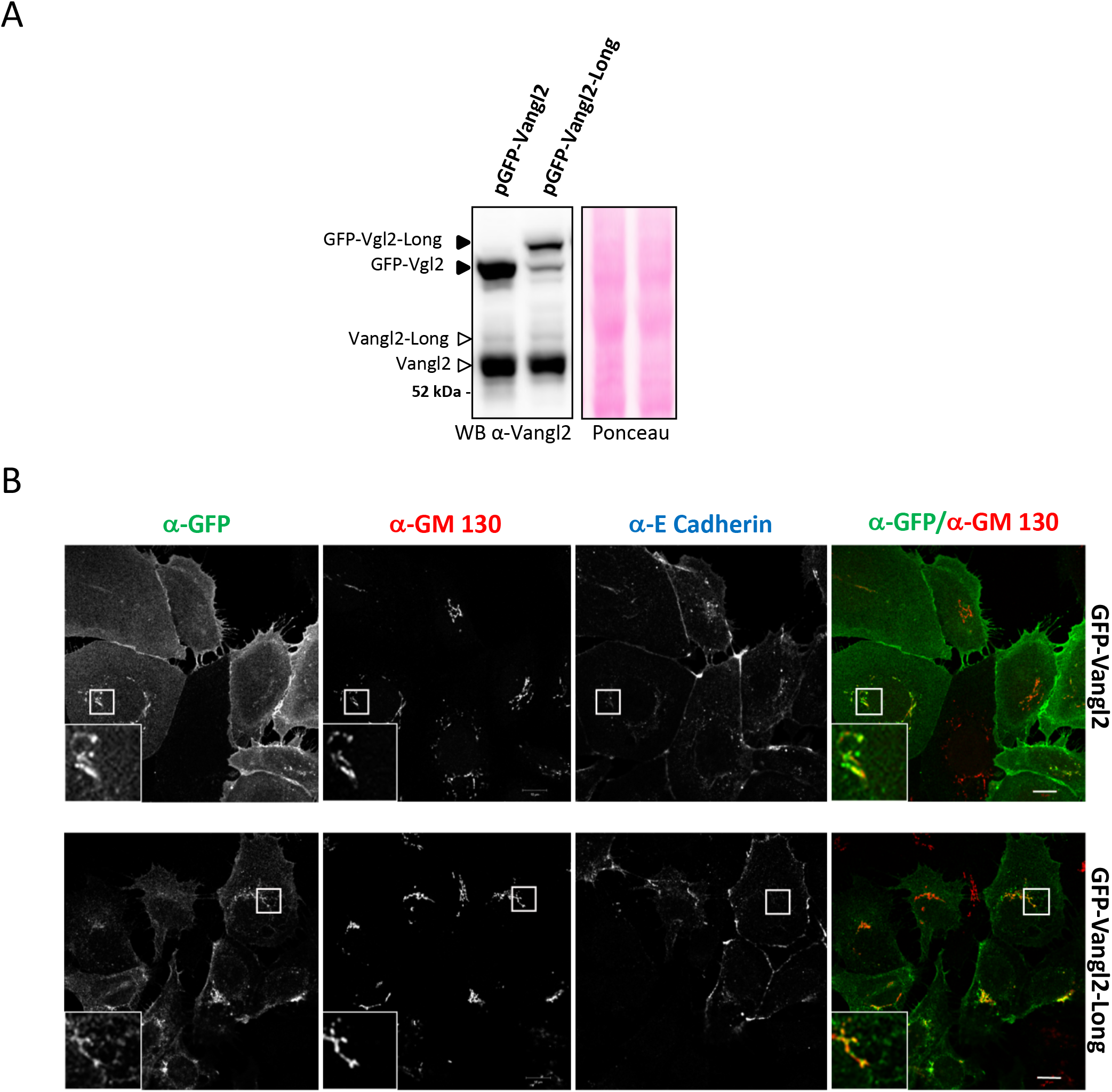
Golgi localization of Vangl2-Long in non-confluent IMCD3 cells. **(A)** Western blot analysis of IMCD3 cells stably transfected with pGFP-Vangl2 or pGFP-Vangl2-Long. Endogenous (open arrows) as well as GFP-tagged (filled arrows) Vangl2 and Vangl2-Long proteins are detected using mAb 36E3. **(B)** The same cells as used in (A) were grown under non-confluent conditions and analyzed by immunofluorescence and confocal microscopy using the indicated. Scale bars correspond to 10 μm.

**Figure S8.**
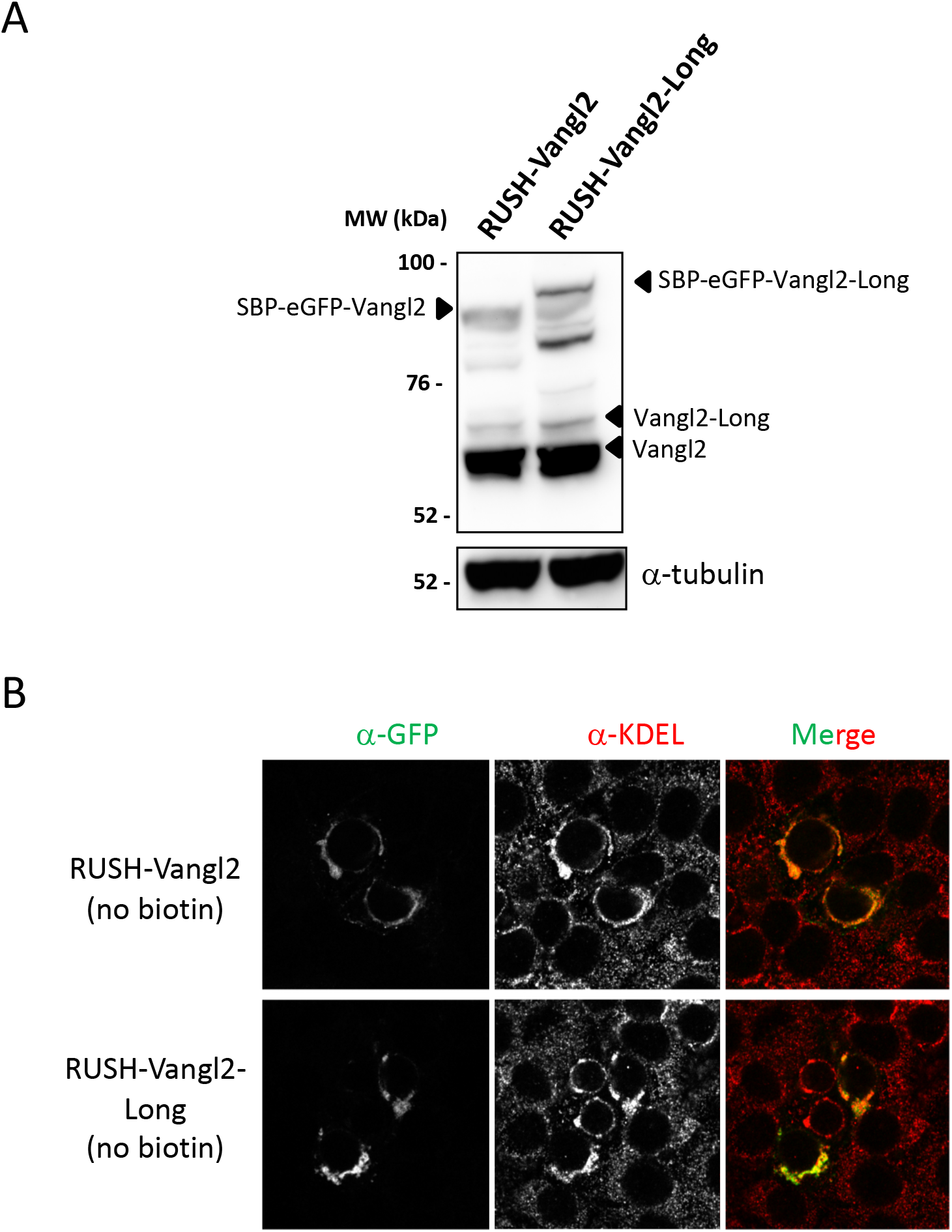
Biochemical and cytological validation of the GFP-Vangl2 and GFP-Vangl2-Long RUSH reporters. **(A)** Western blot analysis of IMCD3 cells transfected with either RUSH-Vangl2 or RUSH-Vangl2-Long plasmids using mAb 36E3 (upper panel) and anti-α tubulin (lower panel). **(B)** Same cells as in (A) and grown in the absence of biotin were analyzed by immunofluorescence and confocal microscopy with a GFP antibody (left panels) or a KDAEL antibody that stains the endoplasmic reticulum. Bar represents 10 μm.

**Figure S9.**
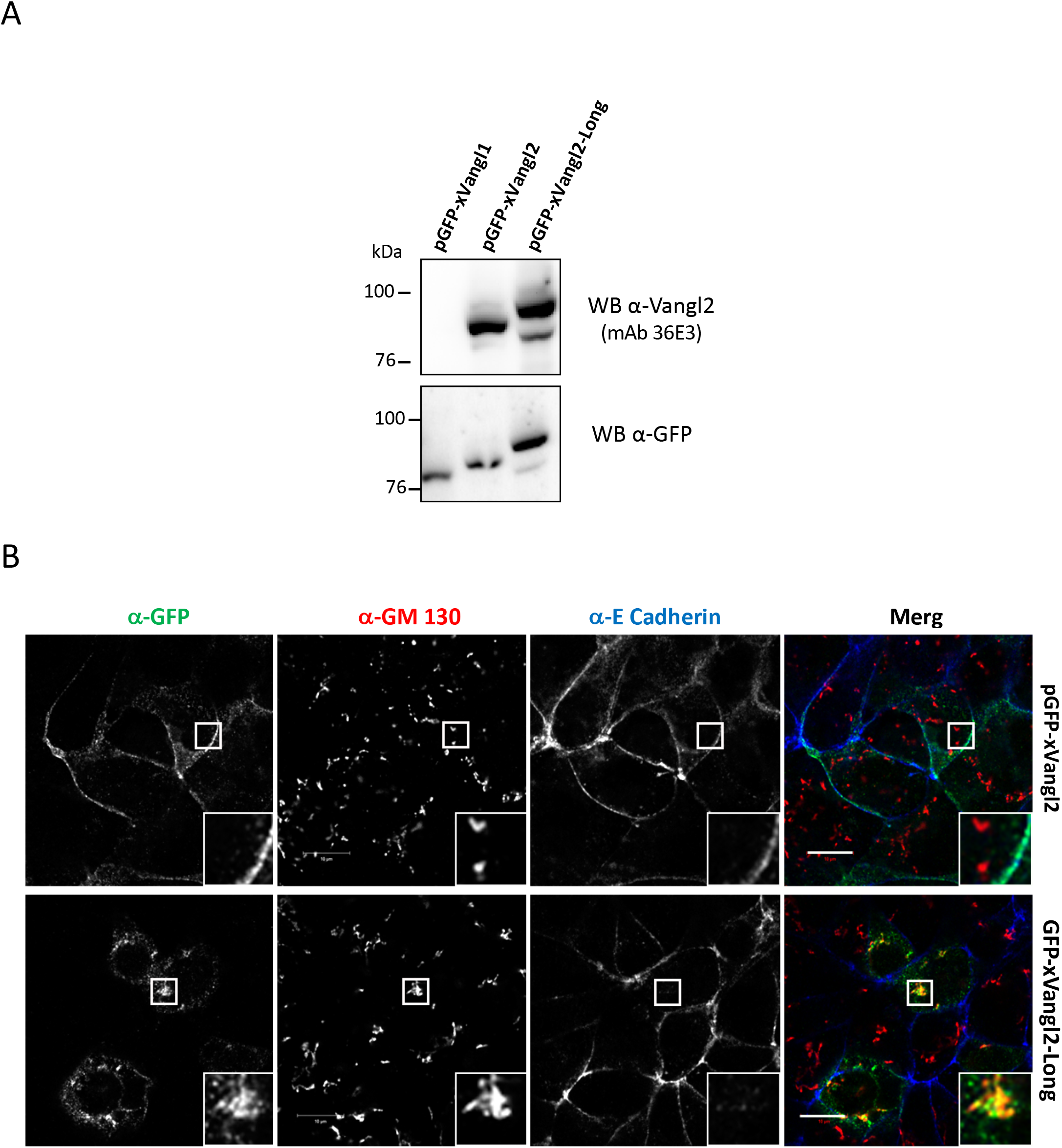
Expression and localization of GFP-tagged *Xenopus* Vangl2 and Vangl2-Long proteins in IMCD3 cells. **(A)** IMCD3 cells transfected with GFP-tagged *Xenopus* Vangl1 (GFP-xVangl1), Vangl2 (GFP-xVangl2) and Vangl2-Long (GFP-xVangl2-Long) constructs were probed by Western blotting with mAb 36E3 (upper panel) or anti-GFP antibodies (lower panel). Note the specific immunoreactivity of mAb 36E3 against the GFP fusions of xVangl2 and xVangl2-Long proteins but not xVangl1. **(B)** IMCD3 cells stably expressing xVangl2 or xVangl2-Long proteins both fused to GFP, were analyzed by immunofluorescence microscopy using antibodies against GFP, E-cadherin and GM130. As observed for the human Vangl2 and Vangl2-long isoforms (see fig. 3), xVangl2 is mostly detected at the plasma membrane (stained with E-cadherin Ab) whereas a large pool of xVangl2-Long is also found at the Golgi apparatus (stained with GM130 Ab).

**Figure S10.**
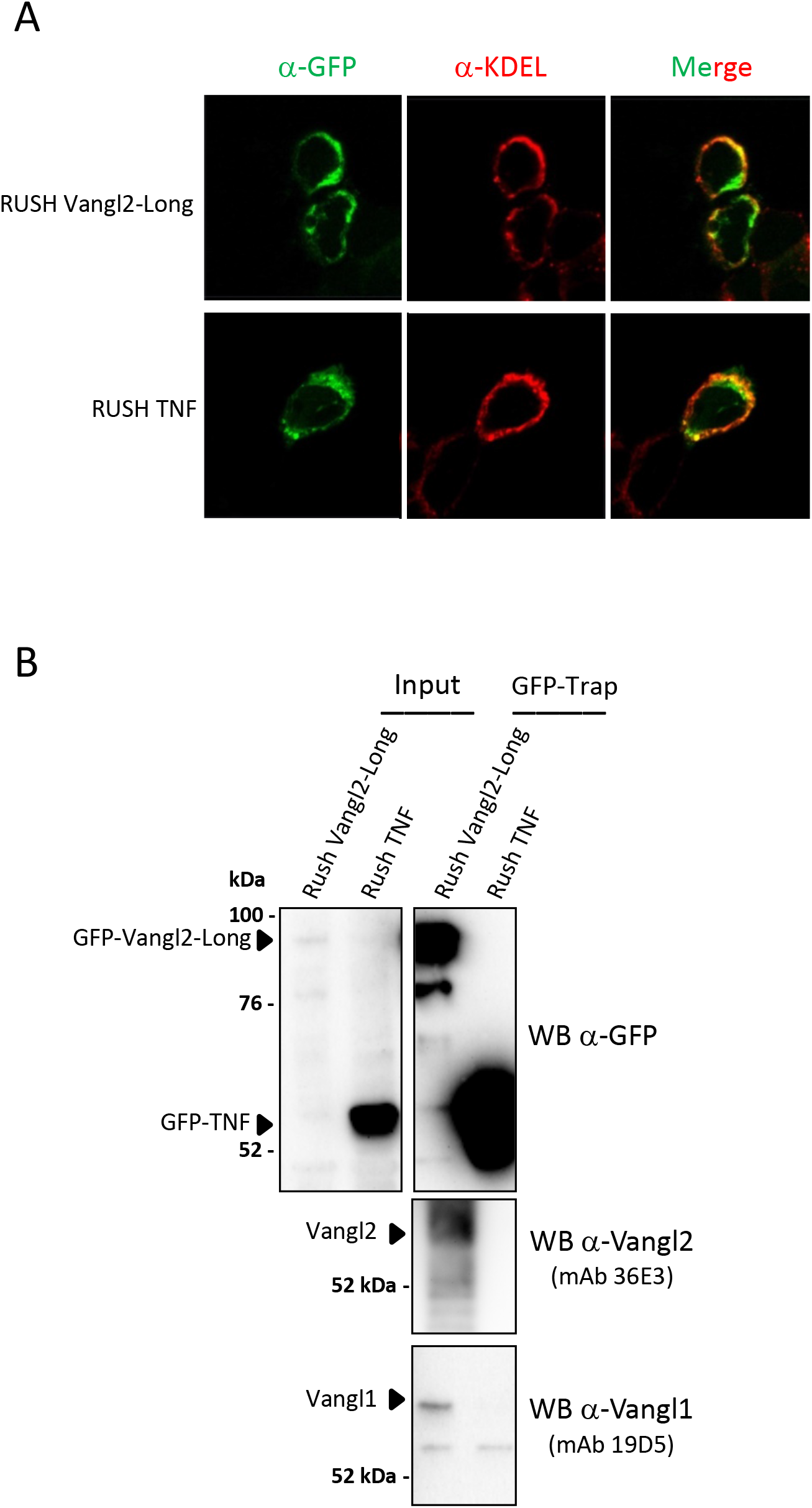
Vangl1 and Vangl2 co-purify with ER-sequestered Vangl2-Long. **(A)** HEK 293T cells were transiently transfected with plasmids encoding an ER-anchored GFP-Vangl2-Long RUSH reporter as described in Fig. 4 or a control ER-anchored GFP-TNF RUSH reporter construct. Two days after transfection, cells grown in the absence of biotin were analyzed by immunofluorescence microscopy using the indicated antibodies. **(B)** Cell lysates from cells prepared as described in (A) were subjected to a GFP-Trap immunoprecipitation prior to Western blot analyses with the indicated antibodies. Note the ability of endogenous Vangl1 and Vangl2 to copurify with the GFP-Vangl2-Long RUSH reporter under conditions where the latter protein is sequestered in the ER in the absence of biotin.

**Figure S11:**
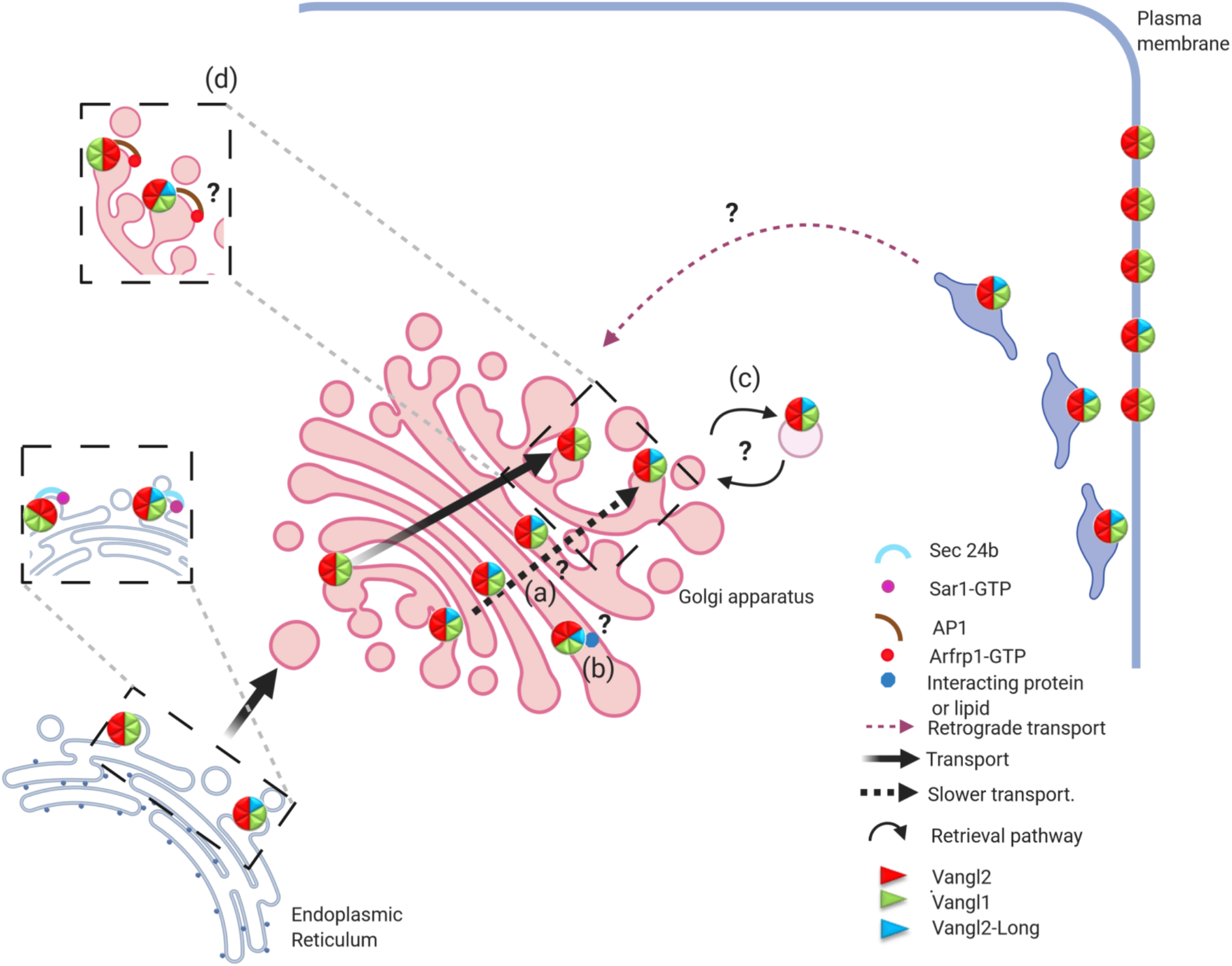
Vangl1 and Vangl2 co-purify with ER-sequestered Vangl2-Long. Model of Vangl2 isoforms specific trafficking from the ER to the plasma membrane and hypotheses. The diagram recapitulates cytological data obtained in the present study with the newly identified Vangl2-Long translational variant that is shown to form tripartite complexes with Vangl1 and Vangl2. We opted for a scenario whereby one Vangl2-Long molecule assembles with 3 molecules of Vangl2 and two molecules of Vangl1, based on the biochemical analysis presented here. A second complex made of three molecules of VAngl1 and Vangl2 is also depicted.

**Table S1:**
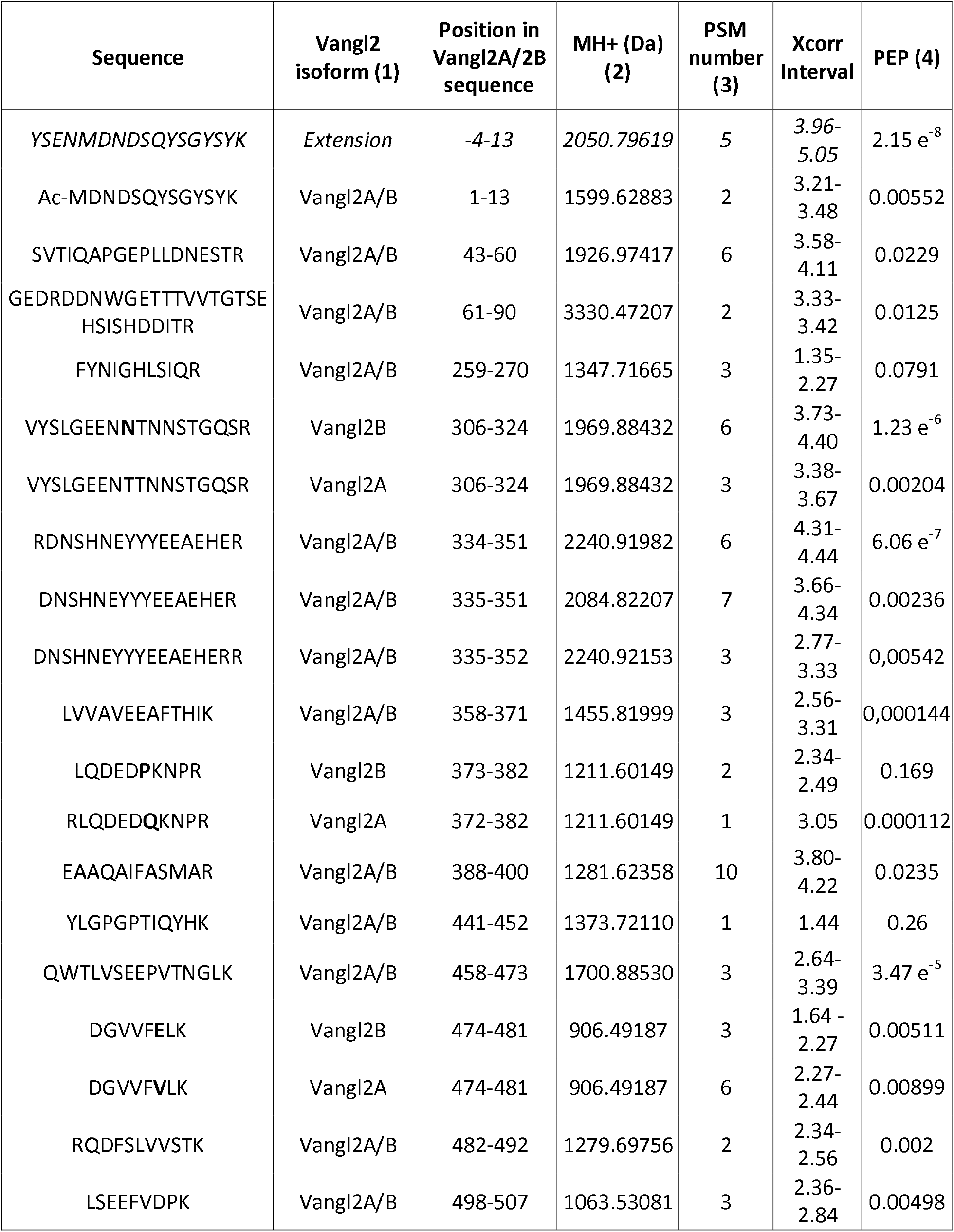
List of Vangl2 peptides detected by LC-MS/MS in the material immunoprecipitated from *Xenopus* embryos with the Vangl2-specific mAb 36E3 antibody. (1) Vangl2 A and B are two isoforms encoded by the *Xenopus laevis* Vangl2 loci (2) MH+ correspond to the monoisotopic masse of monocharged peptide; (3) PSM : total number of identified peptide sequences (peptide spectrum matches) for the protein, including those redundantly identified ; (4) PEP : posterior error probability is the probability that the observed PSM is incorrect.

**Table S2.** LC-MS/MS mass spectrometry analysis of proteins co-immunoprecipitating with xVangl2. (1) TOP3 : calculated as the mean of the three highest peptides areas measured for each protein ; (2) #peptides : number of peptide sequences unique to a protein group. (3) PSM : total number of identified peptide sequences (peptide spectrum matches) for the protein, including those redundantly identified ; (4) Vangl2 A and B are two isoforms encoded by the Xenopus *laevis* Vangl2 loci.

| Run1 |  |  |  |  |  |  |
| --- | --- | --- | --- | --- | --- | --- |
| Accession | Description | TOP3<br>AREA (1) | Sequest<br>Score | Sequence<br>Coverage | #<br>Peptides<br>(2) | # PSM<br>(3) |
| Q90Z05/Q90X64 | Vang-like protein<br>2-A/2-B (4) | 9,379E7 | 62,20 | 35,70 | 15 | 22 |
| A0A1L8H5Y3 | Vang-like protein 1 | 2,567E6 | 10,40 | 10,74 | 4 | 4 |
| Run2 |  |  |  |  |  |  |
| Accession | Description | TOP3<br>AREA | Sequest<br>Score | Sequence<br>Coverage | #<br>Peptides | # PSM |
| Q90Z05/Q90X64 | Vang-like protein<br>2-A/2-B | 1,331E8 | 70,65 | 38,00 | 16 | 23 |
| A0A1L8H5Y3 | Vang-like protein 1 | 2,833E6 | 10,71 | 10,74 | 4 | 4 |
| Run3 |  |  |  |  |  |  |
| Accession | Description | TOP3<br>AREA | Sequest<br>Score | Sequence<br>Coverage | #<br>Peptides | # PSM |
| Q90Z05/Q90X64 | Vang-like protein<br>2-A/2-B | 8,495E7 | 51,62 | 23,42 | 11 | 17 |
| A0A1L8H5Y3 | Vang-like protein 1 | 2,961E6 | 10,91 | 10,74 | 4 | 4 |

**Table S3:** plasmids constructs used in this study.

| **Plasmid name** | **insert** | **origin** | **cloning method** |
| --- | --- | --- | --- |
| pVangl2 | human Vangl2 (aa 1-521) ORF | this study | Gateway |
| pVangl2-SL | human Vangl2 (aa 1-521) ORF + 342bp 5'UTR | this study | Gateway |
| pVangl2-Long | human Vangl2 (aa 1-521) ORF + 144bp 5'UTR (ATA<sup>-144</sup> -> ATG and ATG<sup>1</sup> -> GCA) | this study | Gateway |
| pGFP-Vangl2 | human Vangl2 (aa 1-521) ORF cloned downstream of GFP | Belotti et al., 2012 | Gateway |
| pGFP-Vangl2-Long | human Vangl2 (aa 1-521) ORF + 144bp 5'UTR cloned downstream of GFP | this study | Gateway |
| pGFP-Vangl1 | human Vangl1 (aa 1-524) ORF cloned downstream of GFP | Belotti et al., 2012 | Gateway |
| pGFP-Vangl1-Long | human Vangl1 (aa 1-524) ORF + 144bp 5'UTR Vangl2 cloned downstream of GFP | this study | Gateway |
| pVangl2-SL-ATA-to-GCA | human Vangl2 (aa 1-521) ORF + 342bp 5'UTR ATA<sup>-144</sup> -> GCA | this study | Gateway |
| pVangl2-SL-ATG-to-GCA | human Vangl2 (aa 1-521) ORF + 342bp 5'UTR ATG<sup>1</sup> -> GCA | this study | Gateway |
| pGFP-Vangl2-Long-V8A | human Vangl2 (aa 1-521) ORF + 144bp 5'UTR - V8A cloned downstream of GFP | this study | Gateway |
| pGFP-Vangl2-Long-L11A | human Vangl2 (aa 1-521) ORF + 144bp 5'UTR - L11A cloned downstream of GFP | this study | Gateway |
| pGFP-Vangl2-Long-V8A L11A | human Vangl2 (aa 1-521) ORF + 144bp 5'UTR - V8A L11A cloned downstream of GFP | this study | Gateway |
| pGFP-Vangl1 (Xenopus) | Xenopus laevis Vangl1 ORF | J. Wallingford |  |
| pGFP-Vangl2<br>(Xenopus) | Xenopus laevis Vang2 ORF | J. Wallingford |  |
| pRFP-Vangl2 Long<br>(Xenopus) | Xenopus laevis Vang2 ORF + 159bp<br>5'UTR | This study | Not1-Xho1<br>fragment |
| RUSH-Vangl2 | human Vangl2 (aa 1-521) ORF<br>cloned downstream of SBP-GFP<br>+ ER anchor(Str-Ii) | this study | Fse1-Xho1<br>fragment |
| RUSH-Vangl2-Long | human Vangl2 (aa 1-521) ORF + 144bp<br>5'UTR cloned downstream of SBP-GFP<br>+ ER anchor(Str-Ii) | this study | Fse1-Xho1<br>fragment |
| RUSH-TNF | TNF-SBP-EGFP + ER anchor(Str-KDEL) | F. Perez |  |

## Notes

### Competing Interest Statement

The authors have declared no competing interest.

